# High resolution cryo-EM structures of two potently SARS-CoV-2 neutralizing monoclonal antibodies of same donor origin that vary in neutralizing Omicron variants

**DOI:** 10.1101/2022.12.03.518949

**Authors:** Clayton Fernando Rencilin, Mohammad Yousuf Ansari, Arnab Chatterjee, Suprit Deshpande, Sohini Mukherjee, Randhir Singh, Sowrabha Jayatheertha, Poorvi M. Reddy, Payel Das, Nitin Hingankar, Deepak Rathore, Raghavan Varadarajan, Jayanta Bhattacharya, Somnath Dutta

**Author notes:** equal contribution (Clayton Fernando Rencilin, Mohammad Yousuf Ansari, Arnab Chatterjee). **Corresponding authors**: Somnath Dutta, Jayanta Bhattacharya ( &).

## Abstract

While vaccines have by large been found to effective against the evolving SARS-CoV-2 variants, the profound and rapid effectivity of monoclonal antibodies (mAbs) in significantly reducing hospitalization to severe disease outcomes have also been demonstrated. In the present study, by high resolution cryo-electron microscopy (cryo-EM), we examined the structural insights of two trimeric spike (S) protein bound mAbs isolated from an Indian convalescent individual infected with ancestral SARS-CoV-2 which we recently reported to potently neutralize SARS-CoV-2 from its ancestral form through highly virulent Delta form however different in their ability to neutralize Omicron variants. Our findings showed binding and conformational heterogeneities of both the mAbs (THSC20.HVTR04 and THSC20.HVTR26) bound to S trimer in its apo and hACE-2 bound forms. Additionally, cryo-EM resolved structure assisted modeling highlighted key residues associated with the ability of these two mAbs to neutralize Omicron variants. Our findings highlighted key interacting features modulating antigen-antibody interacting that can further aid in structure guided antibody engineering to enhance their breadth and potency.

**Highlights:**

- Two potent human mAbs obtained from a single donor differ binding to Omicron spikes
- Pattern of binding and conformation of these mAbs bound to full length spike differs
- Antibody binding alters the conformational states of S trimer in its apo and hACE-2 bound forms.
- Cryo-EM structure guided modeling highlighted correlates of interacting residues associated with resistance and sensitivity of BA.1, BA.2, BA.4/BA.5 resistance and sensitivity against these mAbs.

## Introduction

Globally severe acute respiratory syndrome coronavirus 2 (SARS-CoV-2) has infected over 500 million people with more than 6 million deaths (https://covid19.who.int/). Acquisition of few of several key mutations particularly within the spike protein (S) through natural selection towards maintaining virus fitness associated with its continued evolution has resulted in giving rise to variants of concerns (VOCs) (*1*) (https://www.cdc.gov/coronavirus/2019-ncov/variants/variant-classifications.html). Majority of these VOCs responsible for successive waves of infections globally have been found to be associated with increased transmissibility, infectivity, and immune escape (*2–6*). The Omicron variant which succeeded Delta in late 2021 to become globally dominant form has since then have given rise to several sub lineages from BA.1 to the rapidly emerging BA.5 form (2021). (*7–9*). Of these BA.2 became globally prevalent subsequently since March 2021 of the SARS-CoV-2 including India (https://cov.lanl.gov/components/sequence/COV/embers.comp) (*10, 11*) and very recently Omicron BA.4 and BA.5 with identical spike (S) sequence have started to dominate (https://cov.lanl.gov/components/sequence/COV/embers.comp) (*12*).

The trimeric spike (S) of SARS-CoV-2 is a homotrimeric type I transmembrane fusion glycoprotein which facilitates the initiation of the virus infection by engagement of its receptor binding domain (RBD) with the human angiotensin converting enzyme 2 (ACE2) receptor expressed on epithelial cells (*13*). The S protein consists of two subunits, S1 that contains N-terminal domain (NTD) as well RBD required for attachment to host ACE2 receptor and also the major target of neutralizing antibodies, the S2 subunit is involved in membrane fusion. The receptor binding motif (RBM) within S1/RBD region is responsible for extensive binding with the N-terminal alpha helix on the ACE2 receptor (*14, 15*). The S1/RBD-ACE2 binding allows the S2 undergo conformational changes that further the fusion of viral and host membranes to enable the viral RNA to enter into the host cell towards establishing the infection (*16*). The spike trimer exists in an equilibrium between closed and open form based on the engagement with the receptor (*14*). Neutralizing antibodies have been reported to broadly target three distinct sites in the S1 subunit: (a) those which target the NTD region (*17*, *18*), (b) ones which target the ACE2 binding surface of the RBD and RBM and those block S-ACE2 interaction (*2, 19*) and (c) rare ones that target N-linked glycans in RBD without any influence on S-ACE2 interaction (*20*), however demonstrated destabilizing effect of the trimeric S protein (*21–23*).

While booster doses of different vaccines have been shown to be moderately effective against Omicron and its variants, a considerable reduction in the neutralization titers have been reported recently (*24–29*). In addition to vaccine induced antibody responses, majority of the clinically approved therapeutic monoclonal antibodies (mAbs) have also been found ineffective or variably effective against emerging Omicron variants due to accumulation of variety of spike mutations (*24*, *29*–*32*).

Recently, we reported isolation of five SARS-CoV-2 RBD-reactive mAbs from an unvaccinated convalescent donor (C-03-0020) (*33*), of which two (THSC20.HVTR04 & THSC20.HVTR26) demonstrated potent neutralization of SARS-CoV-2 variants of concern; VOC (Alpha, Beta, Gamma and Delta) and other variants of interest; VOI such as Kappa, Delta Plus and those differed in their epitope specificities; however only THSC20.HVTR26 was found to neutralize Omicron B.1.1.529 (BA. 1). On the other hand we recently found that THSC20.HVTR04, while incapable of neutralize BA. 1 variant, it can potently neutralize Omicron BA.2 and BA.4/BA.5 variants (https://doi.org/10.1101/2022.10.19.512979). Using high resolution cryo-EM based we examined the structural insights and conformational flexibilities of the two mAbs when complexed with the full-length S trimer at 4.5 Å in its pre-fusion form. Using the high resolution cryo EM structure and molecular modeling, we not only determined the detailed interactions between epitopes within spike RBD with the paratopes of both the mAbs (examined as fragment antigen binding regions or Fabs) but also determined the determinants associated with sensitivity and resistance both the mAbs against Omicron BA.1, BA.2 and BA.4/5 variants.

## Results

### Broadly neutralizing THSC20.HVTR04 mAb demonstrated strong binding to Omicron BA.2 and BA.4/BA.5 spikes expressed on cell surface and displayed favorable physicochemical properties

Previously, we reported isolation of five RBD-reactive neutralizing mAbs from an unvaccinated convalescent donor (C-03-0020), two (THSC20.HVTR04 & THSC20.HVTR26) of which demonstrated potent neutralization of SARS-CoV-2 variants of concern; VOC (Alpha, Beta, Gamma and Delta) and other variants of interest; VOI such as Kappa, Delta Plus and also found to show protective efficacy at low dose in K18 hACE2 transgenic mice^1^. Of this two, THSC20.HVTR04 while found to retain its neutralization activity against the Omicron BA.2 and BA.4/BA.5, THSC20.HVT26 could only neutralize BA.1 but not the others. In the present study, we further assessed the ability of these two mAbs to bind to the currently circulating and rapidly emerging Omicron BA.2 and BA.4 / BA.5 variants of concern spikes by carrying out and cell surface binding. As shown in Figure 1A, THSC20.HVTR04 demonstrated strong binding to the spikes of BA.2 and BA.4 expressed on the surface of 293T cells in a dose-dependent manner, while THSC20.HVTR26 did not show any measurable binding to BA.2 and BA.4 spikes expressed on cell surface (data not shown). The dose-dependent binding of THSC20.HVTR04 to the Omicron BA.2 and BA.4/BA.5 variants expressed on cell surface indicating their ability to bind to trimeric spike proteins of BA.2 and BA.4 in its prefusion state. When assessed for their physicochemical properties, both THSC20.HVTR04 and THSC20.HVTR26 were found to be highly thermostable up to 70°C as assessed by differential scanning calorimetry (DSC) (Figure S1C), demonstrated excellent homogeneity (Figure S1B) by dynamic light scattering (DLS). The correlation coefficient function showed single decay component. The size distribution by intensity showed a single effective peak of diameter around 12.93nm and 13.57nm respectively, with negligible evidence of aggregates or other components. The data suggest that the purified mAbs are homogeneous and no IgG aggregation observed when we assessed their SEC profiles (Figure S1A). These findings validate the favorable developability profiles of both the mAbs for clinical development.

**Fig 1.**
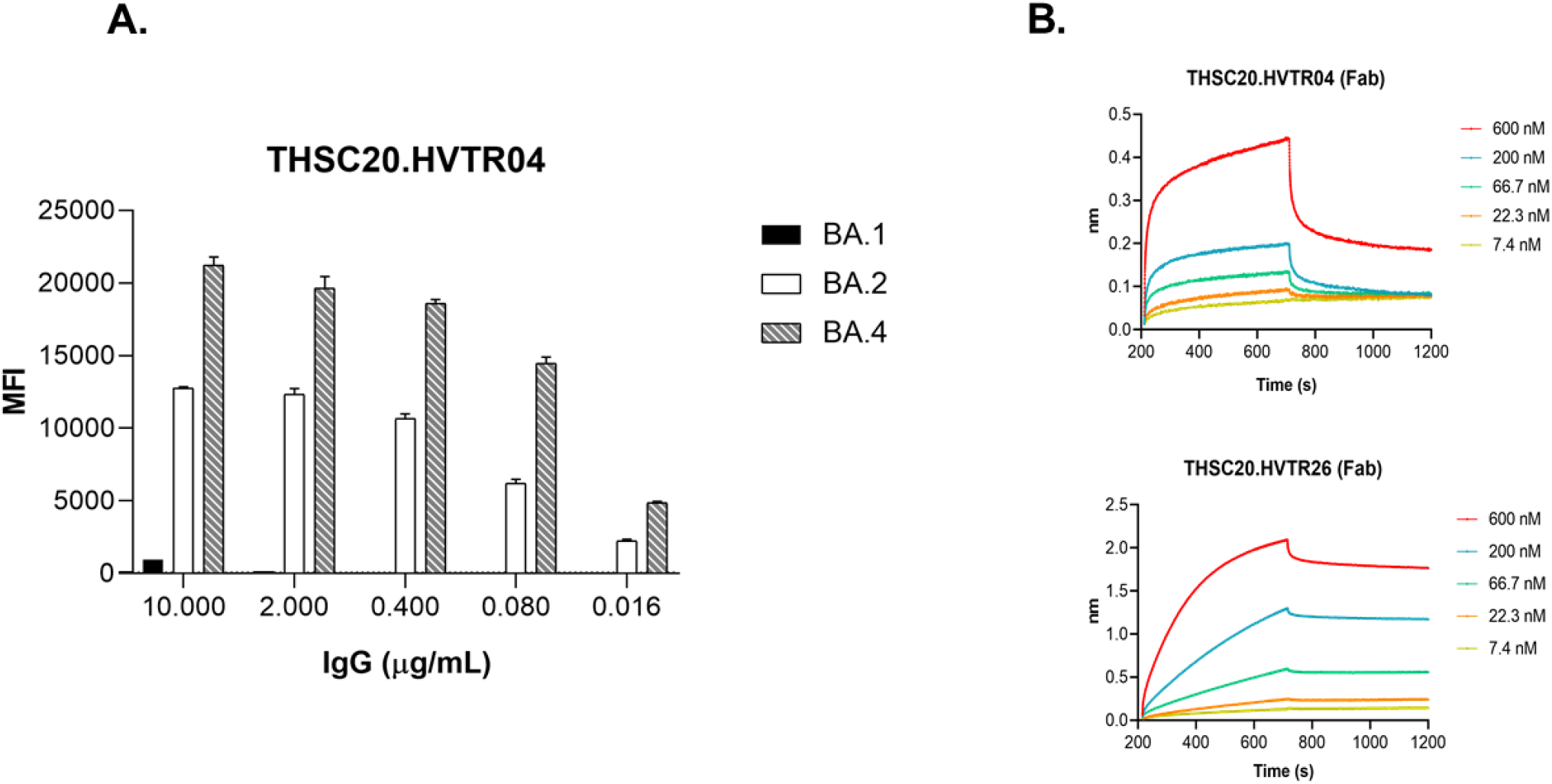
Binding of Omicron BA.2 and BA.4/BA.5 variants to THSC20.HVTR04 and THSC20.HVTR26 mAbs. A. Cell surface binding assay B. the binding kinetics of Fab4 and Fab26 to SARS-CoV-2 RBD protein.

### Structural properties of binding modes of S protein bound Fab4 and Fab26 complexes by cryoEM analysis

We first examined the binding potential of the Fabs prepared from THSC20.HVTR04 and THSC20.HVTR26 Fabs to the SARS-CoV-2 RBD by BLI (Bio-layer interferometry) before proceeding to examine their structural insights by TEM (Transmission Electron Microscopy) study. Both the purified mAb4 and mAb26 Fabs (referred to as Fab4 and Fab26 from hereon) were found to bind to RBD (**Figure 1B**) ensuring that they are fit to proceed for EM experiments. Interestingly, mAb4 Fab demonstrated a faster on-rate and a faster off-rate compared to mAb26 Fab indicating that Fab4 requires less energy for interacting with the SARS-CoV-2 spike protein than Fab26. On the other hand, slow off-rate displayed by Fab26 indicated that its interaction with RBD spike is more stable than that of Fab4. Next, we carried out negative stain TEM analysis to first predict the frequency of Fab bound forms of spike protein complexes. SARS-CoV-2 spike (S) protein was incubated with the two Fabs (Fab4 and Fab26) for TEM imaging. Negative staining (NS) TEM imaging and reference-free 2D classifications were performed to investigate the sample quality and to visualize the dynamic nature of spike-antibody interactions. Our TEM studies showed that the majority of the S proteins were highly homogeneous triangular cone-shaped molecules with some extra densities attached with RBDs of S2 region (Figure S2), which was found to be completely absent in previous negative staining images of only spike protein at different pH (*15*). Additionally, no tendency of aggregation in spike protein was observed in the presence of Fab4 and Fab26s (Figure S2). Reference-free 2D class averages of spike protein with Fabs indicated that extra densities were attached to the RBDs. Additionally, it appeared that RBD is in up conformation when it binds to Fab4 and Fab26. In 2D class averages, only one Fab, two Fab, and three Fab densities were clearly visible in different class averages, which agrees with the recently published data of the same antibodies with Fab fragments (*33*). Overall, the NS TEM analysis permitted us to predict the frequency of Fab-bound forms of spike protein complexes (Figure S2).

Next, through the construction of cryo-EM images at high resolution, we examined the morphological changes of S protein protomers with both the Fab complexed with S trimer under physiological conditions. The Fab fragments of THSC20.HVTR04 (Fab4) and THSC20.HVTR26 (Fab26) were incubated with SARS-CoV-2 spike ectodomain and imaged at cryogenic temperature for analyzing their structural properties. Previous studies indicated that S protein was subdivided into S1 and S2 subdomain, where RBD and NTD were located in S1 subdomain. In our current study, we were able to see extra densities at RBDs in the S2 region of S protein in cryo-EM images and reference-free 2D class averages. (Figure S3). Majority of the 2D class averages indicated that the S protein either interacts with two Fabs or three Fab complexes, and this phenomenon was noticeable for both the Fabs (Figure S3A, B). Therefore, 3D classification and refinement were performed to characterize the high-resolution 3D structure of S protein bound to Fab4 and Fab26 (Figure S4, S5 and S7). For both the data sets, low resolution (40Å) spike trimer was first used as an initial model (EMDB) where Fab density appeared near RBD region after 3D classification (Figures S4 and S5). The Fab densities were clearly visible and firmly connected with RBDs, which were supported by our negative staining TEM and BLI data. Furthermore, 3D classification techniques were applied to classify the images into various classes, where the structural variabilities of RBDs were clearly visible. Furthermore, we were able to determine the high-resolution cryo-EM maps of Fab4 complexes, having global resolutions of 4.54, 5.15, and 4.90 Å (Figure 2A-C, 5.A-F, Figure S6) respectively. In total, we found the Fab4-S trimer complex in three different states, i.e., two Fabs bind to S protomer (state I-initial stage), three Fab binding to S protein (State II – intermediate stage and State III – final stage) (Figure 2A-C, 5A-F). Detailed interactions between Fab4-S protein was performed, which discussed later of this study. Moreover, we were able to identify the actual Fab-RBD binding propensity from this 3D classification studies which indicated that Fab4 adopts two Fab and three Fab binding conformations. However, the lesser proportion of the Fab4-S protein complex was observed to be in three Fab binding states, where three Fabs interact with three RBDs and all three RBDs were in the up and partial open conformation (Figure 2, S4 and S5). A major population also observed to co-exist where two Fab tightly interacted with two up conformations of RBDs, whereas the remaining one RBD having down-close conformation. Similar conformational variabilities were observed in Fab26 with S-protein (Figure 2D, E: 5G-J), where two Fab26 tightly interacted with two up conformation RBDs (Figure 2F-G). These results suggest that both the antibodies strongly interact with spike RBD. However, huge conformational heterogeneities were observed in RBDs and Fab structures. Thus, we targeted to identify conformational variabilities of the complexes.

**Fig 2.**
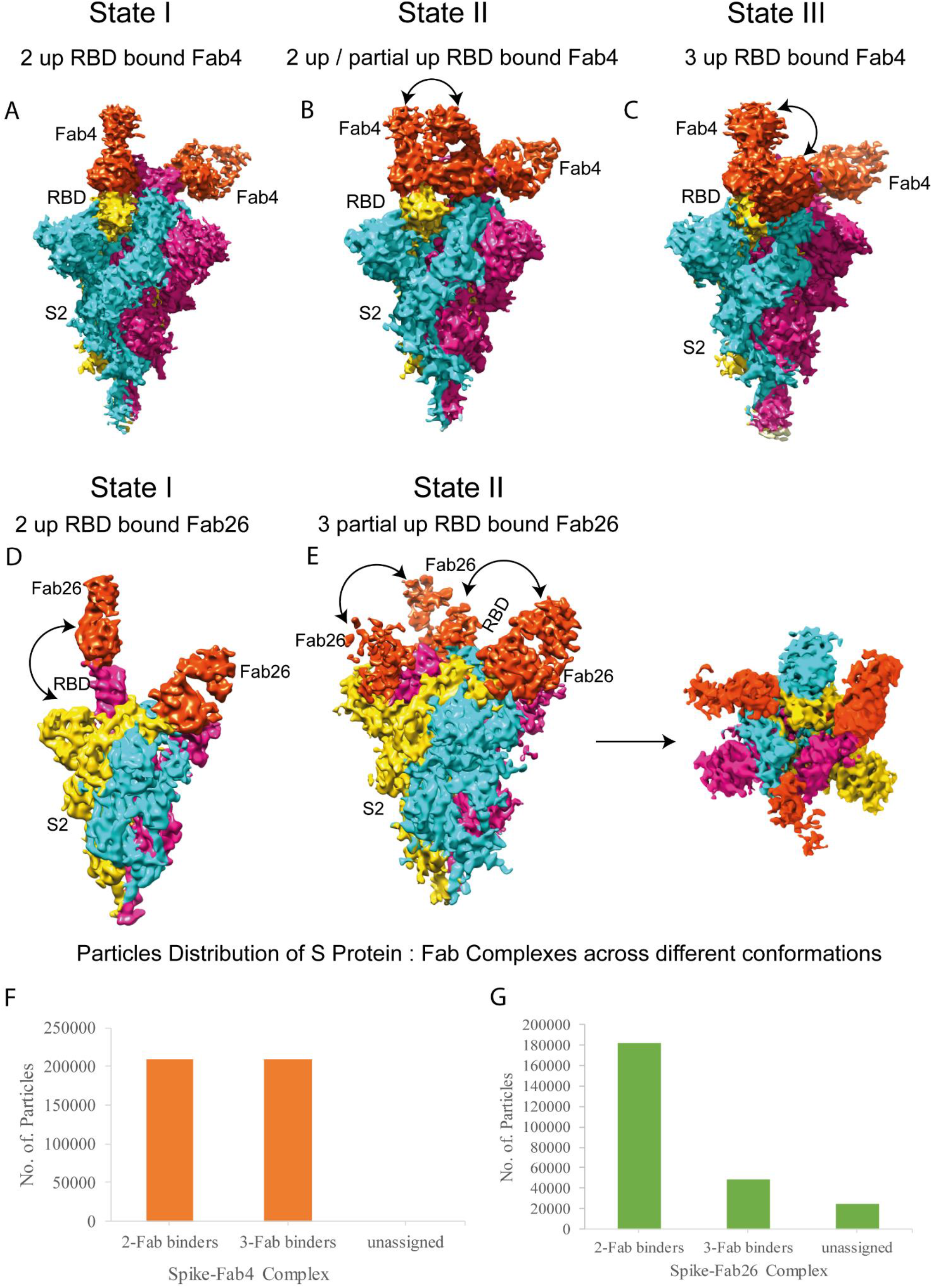
Characterizing different conformational states of Fab4- and Fab26 bound S trimer complexes: A. State I represent a Cryo-EM map two Fab4 bound to two “up” RBD conformation protomers. Shifting of one RBD is shown in arrow while binding to the Fab4. The color of the first, second, and third protomers are gold, magenta, and cyan respectively. Fab is colored in orange red. **B**. State II represents the Cryo-EM of two Fab4 bound to “up” RBD and one Fab4 bound to partial open RBD form. **C**. State III represents an atomic model of three Fab4 bound to a partial open form of the protomers. On the extreme right, perpendicular views of state II is given. Shifting of all the RBDs from each other is shown in the arrow upon binding to Fab4. **D**. State I represent atomic model two Fab26 bound to two “up” RBD conformation protomers. Shifting of one RBD is shown in the arrow while binding to the Fab26. **E**. State II represents an atomic model of three Fab26 bound to the partial open form of the protomers. On the extreme right, perpendicular views of state II are given. Shifting of all the RBD from each other is shown in the arrow upon binding to Fab26. **F, G**. Bar graphs represent the number of particles present in 2-Fab and 3-Fab binding conformations of Fab4 and Fab26 with S protein complexes respectively.

### Conformational dynamics of SARS-CoV-2 S protein complexed with Fab4 and Fab26

We next analyzed the conformational dynamics of the spike bound to Fab4 and Fab26 complexes. At first, we analyzed the Fab4-S trimer complex captured in three different conformations: first fashion with two fab binding and then second with three Fab’s binding, where three Fab binding to the S protein adopts two different conformations (Figure 2 A-C). As mentioned earlier, we determined several cryo-EM maps of spike-Fab complexes at different resolutions. However, we took into account the most high-resolution EM-map of the Fab4-spike complex to analyze the fine details of the Fab4-RBD interaction (Figure 3A, 4A). The available atomic model structure (PDB ID: 7KMS) was next fitted into EM maps, which was further refined in real space refinement to calculate the atomic model for the all the conformational states to identify the detailed conformational changes (Figure 5A-F) in the Fab4-S trimer complexes. This pseuedoatomic model of spike protein with two Fab4 complexes, identifies the precise identity of the epitopes for each protomer from each state represented as heatmap in Figure 3B. A total of 16 amino acid residues were identified interacting with Fab4 and the Fab4 interacting sites are N440, K444, V445, G446, G447, P499, and T500, which agrees with our previous observation (*33*). Fab4 belongs to the family of IGHV3-53 antibodies and based on the interaction studies, we concluded that Fab4 comes under most immunodominant Class1 mAbs targeting RBD (*34*). This data also supports the observation of the inability of THSC20.HVTR04 in neutralizing Omicron BA. 1 (*33*) due to the presence of G446S mutation in the RBD domain. However, rest of the interacting amino acids exist in BA. 1 variant and we anticipated weak interactions between those amino acid residues in BA. 1 spike and THSC20.HVTR04. This weak interaction between BA.1 spike and THSC20.HVTR04 was validated using BLI data. The interfacial area of Fab4 in complex with S trimer was mapped at 729 Å^2^, with 598- and 131 Å^2^ for heavy and light chains respectively. We observed that 11 residues participated in non-hydrogen binding with Fab4. The CDRH3 region of Fab4 heavy chain residues E107 and W108 were found to be involved in interacting with N440, and N437 respectively (Figure 3 E, F) whereas N439 forms H-bond with the residue N31 in the CDRH1 region similar to the CDRH3 interacting residues (Figure 3F). The paratopes Y51, P57, V105, and P106 were found to be involved in hydrophobic contacts with the epitope V445 (Figure 3D). Interestingly, a hydrophobic pocket was found to be created by combination of the three chains, such as light and heavy chains of Fab4 and S protomer. Paratope Y51 and P57 were found residing on the CDRL2 region, while V105 and P106 residues from CDRH3 regions constructs the stable hydrophobic pocket interaction with the epitope (Figure 3D). Besides, two hydrogen bonding pairs were found between the complexes with residue N440 establishing a strong interaction with paratope E107 than others with a distance of 2.69 Å. Simultaneously, the paratope N31 forms a hydrogen bonding with epitope N437 with a distance of 3.20 Å. These structural observations demonstrated that two of the CDRHs (CRDH1 and CDRH3) and one of the light chains (CDRL2) of Fab4 plays a major role in potentially neutralizing SARS-CoV-2 by targeting the S Protein (Figure 3C, E). As mentioned earlier, we observed the three different conformations of Fab4-S trimer complex, i.e., two Fabs bind to S protein (state I-initial stage), three Fab binding to S protein, where two states were visible, state II – intermediate stage and state III – final stage) (Figure 5A-F). At state I, epitopes found to commonly link the complex for both protomers in the same state are G446, G447, K444, N440, P499, T500, and V445. To understand the changes in the binding of Fab’s, for each protomer, the complexes were further superimposed, where we observed the shift in the angle of RBD binding of Fab4 to 16.4° from each other. In the subsequent stage, when the third Fab binding to the free protomer 3 of S protein was examined, a significant number of conformational changes identified between the protomer-Fab4 complexes in all the protomers. The structural similarity of protomer 2 and 3 complexes were found to be very similar to each other, which signifies that all protomer complexes are comparably analogous. However, the angle of protomer 2 and 3 showed a considerable change of 6.9 ° and 9.8 ° with respect to protomer 1 for interaction. Interestingly, we observed the common amino acid residues that form epitopes for THSC20.HVTR04 (K444, P499, S443, T500, and V445) found in protomers-2 and −3 did not lose any interaction with protomer 1. Besides, the epitopes G446, and Y449 were found not to participate in the protomer-3 complex (*14*). Finally, in state III, protomer 2 was found to regain its initial form, while protomer 1 remains in the same conformation. The protomer 3 complex adopts the same conformation as protomer 1, which stabilizes the conformation of the S trimer-Fab complex. Additionally, Fabs interaction with protomers 1 and 2 were found to be identical and the RMSD score of individual protomers between the states were found to be 0.321 Å and 0.372 Å respectively. On the account of the third Fab4 binding in the intermediate stage, severe structural variability happened in the pre-existing protomer complexes. Conformational change of protomer 1 and 2 between states I and II are 0.951 Å and 1.113 Å respectively as calculated by their RMSD scores. In addition, the angle of shift observed for the same is 9° and 3.2°. Besides, protomer 3 shifts to 25.3° to accommodate the third Fab in a stable complex (Figure 5A-F). Overall, 21 amino acid interacting partners were observed on Spike to Fab4 in all the conformations. The pattern of epitopes from promoters 1, and 2 of the stabilized states (I and II) were found to be almost similar with the exclusion of a few residues (L441, N437, N448, N450, Q506, S443) (Figure 3B). The variation in interacting residues accompanying one or two protomers observed could be the effect of third Fab binding in protomer 3 (pro3). The common epitopes involved in the interaction network of S protein-Fab4 complex in the steady states are G446, G447, K444, N440, P499, T500, and V445. In addition, all the epitopes for the third Fab is the same with protomer 1 of state I and II, representing the equivalent pattern of binding (Figure 3B). We found two conformational states of Fab26-S complexes: in state I, we observed an asymmetric S trimer with two “up” RBDs forming a complex each with one Fab (Figure 2D). The state II consists of symmetric trimer bound to three RBDs in the sa me “up” conformation, where each RBD participated to form a complex with each of the Fab26 (Figure 2D, E; 5G-J). We refined state I EM map after masking to achieve a high-resolution map of Fab26 and S protein complex for detailed structural analysis (Figure 4A). After a careful examination of the structural properties of the 4.4 Å map of one Fab26 bound to S protein complex, 11 amino acid residues in spike protein were identified to participate in the interacting network with Fab26 and the overall interacting interface area found was 636 Å^2^ (361 Å^2^ on heavy and 275 Å^2^ on light chain) (Figure 4B, D). The number of interfacial residues involved in interaction with heavy and light chains were found to be 7 and 5 respectively with 11 epitopes, which is likely to be the basis for potent neutralization conferred by THSC20.HVTR26 Fab (Figure 4B). The epitope residue F486 interacts with the hydrophobic pocket formed by paratopes L34, Y93, and G112. The RBD is stabilized by this hydrophobic pocket formed in the region between CDRH3 and CDRL3 (Figure 4E). To pause the flexible movement of the RBD one of the aromatic residues, F486 in the epitope region protrudes towards the pocket formed by the β-sheets formed by the CDRL1, CDRL3, and CDRH3 regions (Figure 4E). Overall, the promising amino acid residues forming epitopes for Fab26 on S protein were found to be T478, N481, V483, E484, F486, C488, and Y489. These residues were observed to frequently participate in the interaction network of S protein-Fab complexes (Figure 4B). The amino acid residues N481, and Y489 of the epitopes form a hydrogen bonding with the paratopes Y32, and A101 respectively from the heavy chain regions (CDHR3) (Figure 4F). Our analysis further revealed that the CDRL1 region residues Y32 and N33 interact with N481 and T478 and S97 of the CDRL3 region interacts with the epitope V483. Other potential paratopes found were Y32, S96, S97, and D110, which were also found to interact with the same loop of epitope region (Figure 4F, G). Taken together, our data indicate that towards the neutralization of SARS-CoV-2, Fab26 utilizes heavy chain CDR (CDRH3) and two of the light chain CDRs (CDRL1, and CDRL3) to stall into the receptor-binding ridge of the S protein (Figure 4D). Similar to Fab4, Fab26 also belongs to the IGHV3-53 antibodies germline and the identification of the potent epitopes suggested that Fab26 belongs to most immunodominant Class1 mAbs targeting RBD. Apart from that, we observed another state of conformation in which the third Fab26 was found to bind to the free RBD in the S protein (*34*). To accommodate the third Fab26 all the protomers adopted partial open conformation of S protein and it seems to be stabilized (Figure 2D, E). The number of two Fab26 bound S trimer population was found to be significantly higher as compared to the three Fab26 bound form (Figure 2G).

**Fig 3.**
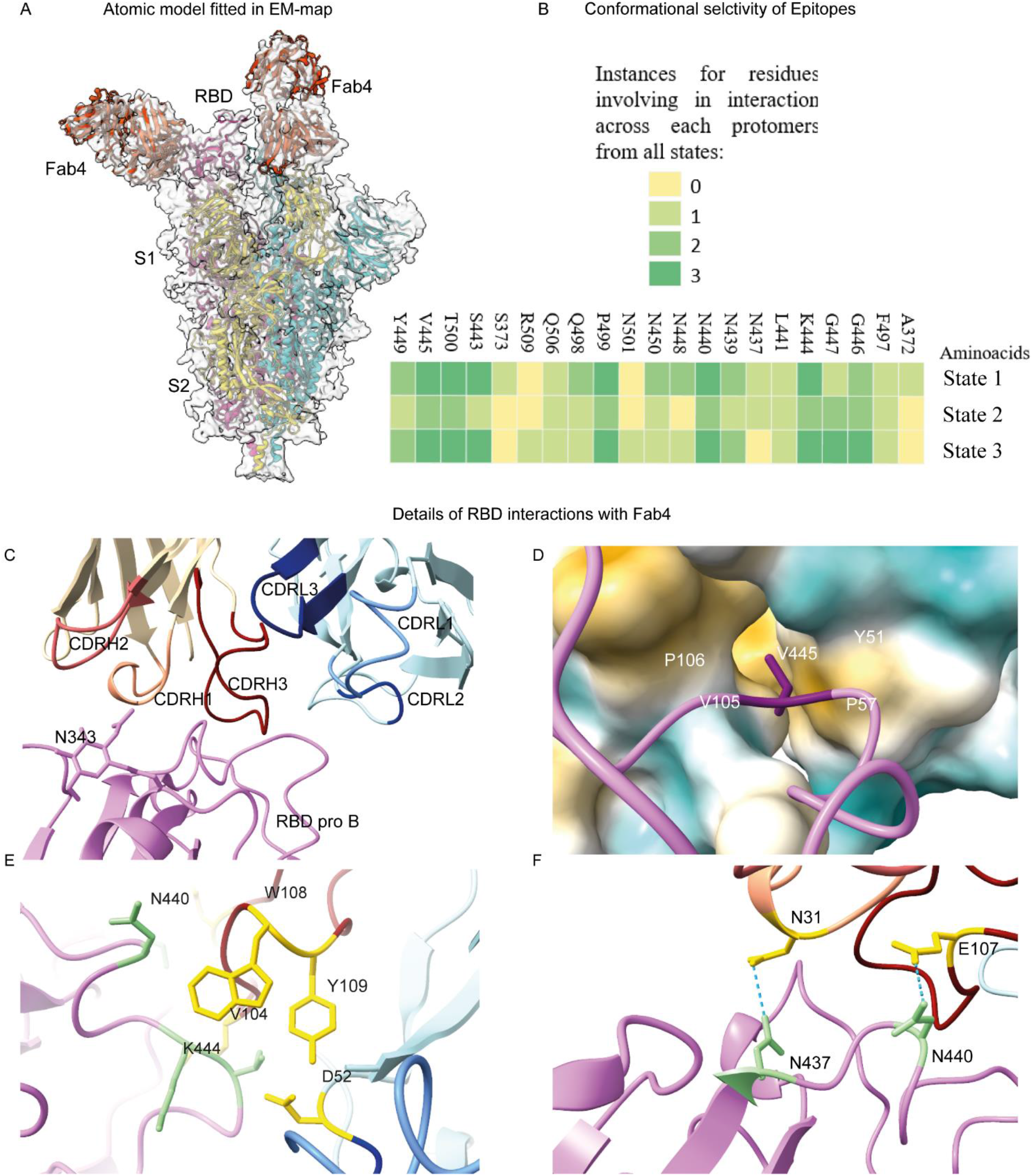
Structural basis of Fab4 accommodation in SARS-CoV2 Spike trimer: **A**. Fitted atomic model generated using Phenix in the high-resolution EM map of S protein-Fab4 complex. The color of the first, second, and third protomers are gold, magenta, and cyan respectively. Fab4 is colored in orange red. **B**. Heat map represents the epitopes residues selectivity and instances for different conformations occurred. It demonstrates the potent epitope residues in all binding modes. **C**. Details of RBD interactions with Fab4. Upper left, extensive hydrophilic interaction on the surface of RBD and Fab4. CDRH1, CDRH2, CDRH3, CDRL1, CDRL2, and CDRL3 are colored as light salmon, Indian Red, Marron, Cornflower blue, Median blue, and Dark blue respectively, and RBD region colored with hot pink. **D**. One residue V445 is shielding in a hydrophobic pocket of Fab4 comprising P57, V105 and P106 residues, **E**. Interaction by hydrogen bonding between Fab4 and RBD and the residues involved in the RBD region such as N437, and N440 are interacting with N31, and E107 respectively. Interacting residues of Fab4 and RBD are colored in gold and pale green respectively. **F**. Interacting partners of Fab26 and RBD involved in non-hydrogen bonding i.e Van-Der-Waals distance.

**Fig 4:**
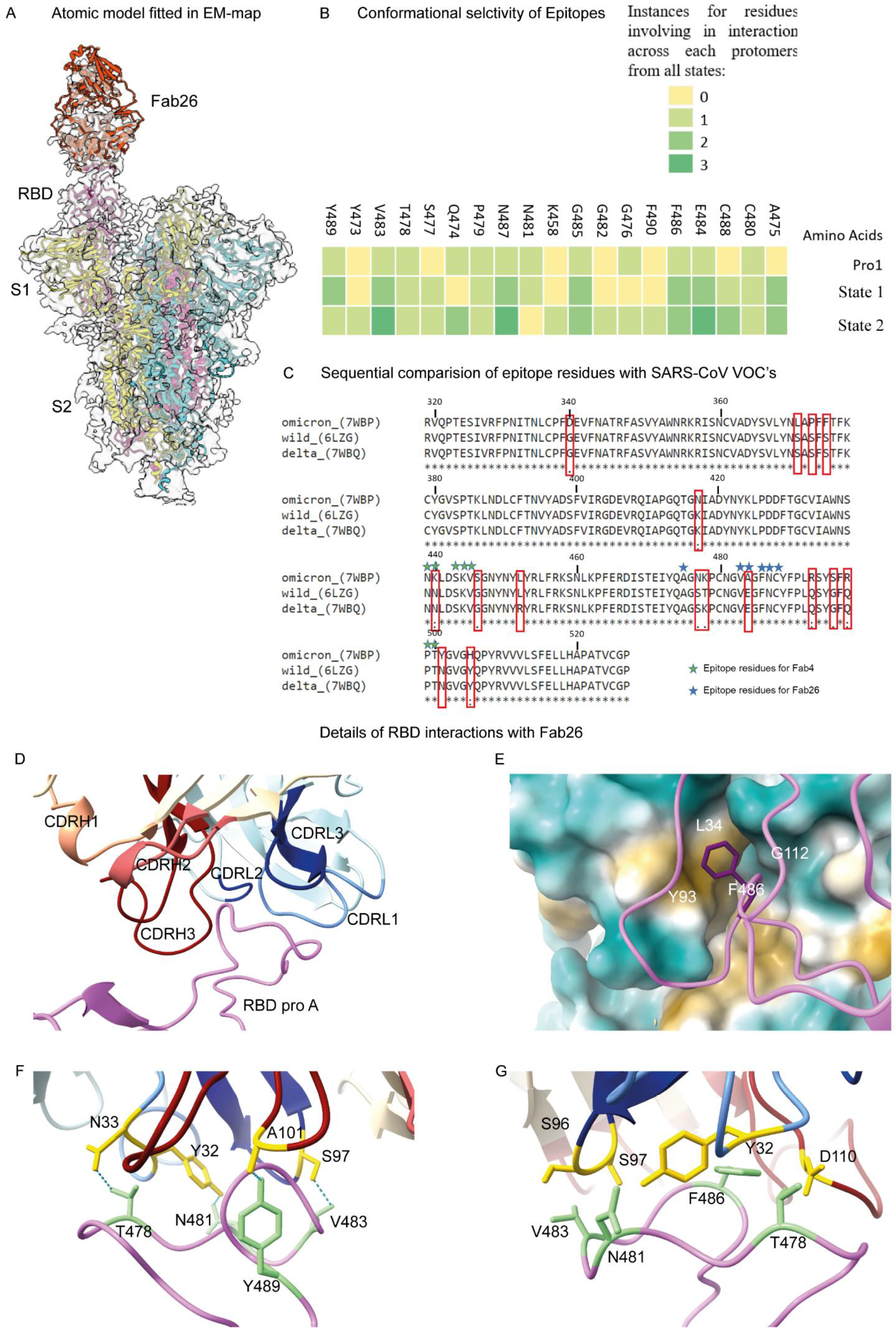
Structural basis of Fab26 accommodation in SARS-CoV2 Spike trimer: **A**. An atomic model was generated using Phenix refinement and fitted in the highest resolution EM map. The color of the first, second, and third protomers are gold, magenta, and cyan respectively. Fab is colored in orange-red. **B**. Characterizing the potent epitopes by comparing the Fab26 binding residues of each protomers across different states. Heatmap showing instances of residues involved in interactions across each protomers from all states and likelihood of these epitopes present in every state of RBD-Fab26 bound forms. **C**. Sequential analysis of Fab4 and Fab26 epitope residues with SARS-CoV2 variants (wild, delta, and omicron). Details of RBD interactions with Fab26. **D**. Extensive hydrophilic interaction on the surface of RBD and Fab26. CDRH1, CDRH2, CDRH3, CDRL1, CDRL2, and CDRL3 are colored as light salmon, Indian Red, Marron, Cornflower blue, Median blue, and Dark blue respectively, and RBD region colored with hot pink. **E**. The common epitope F486 is shielding in a hydrophobic pocket of Fab comprising L34 and G112 residues. **F**. Bottom left is showing hydrogen bonding residues of Fab26 and RBD. Residues in the RBD region such as T478, N481, Y489, and V483 are interacting with N33, Y32, A101, and S97 respectively. Interacting residues of Fab26 and RBD are colored in gold and pale green respectively. **G**. The bottom right shows various interacting partners of Fab26 and RBD in the Van-Der-Waals distance.

**Fig 5:**
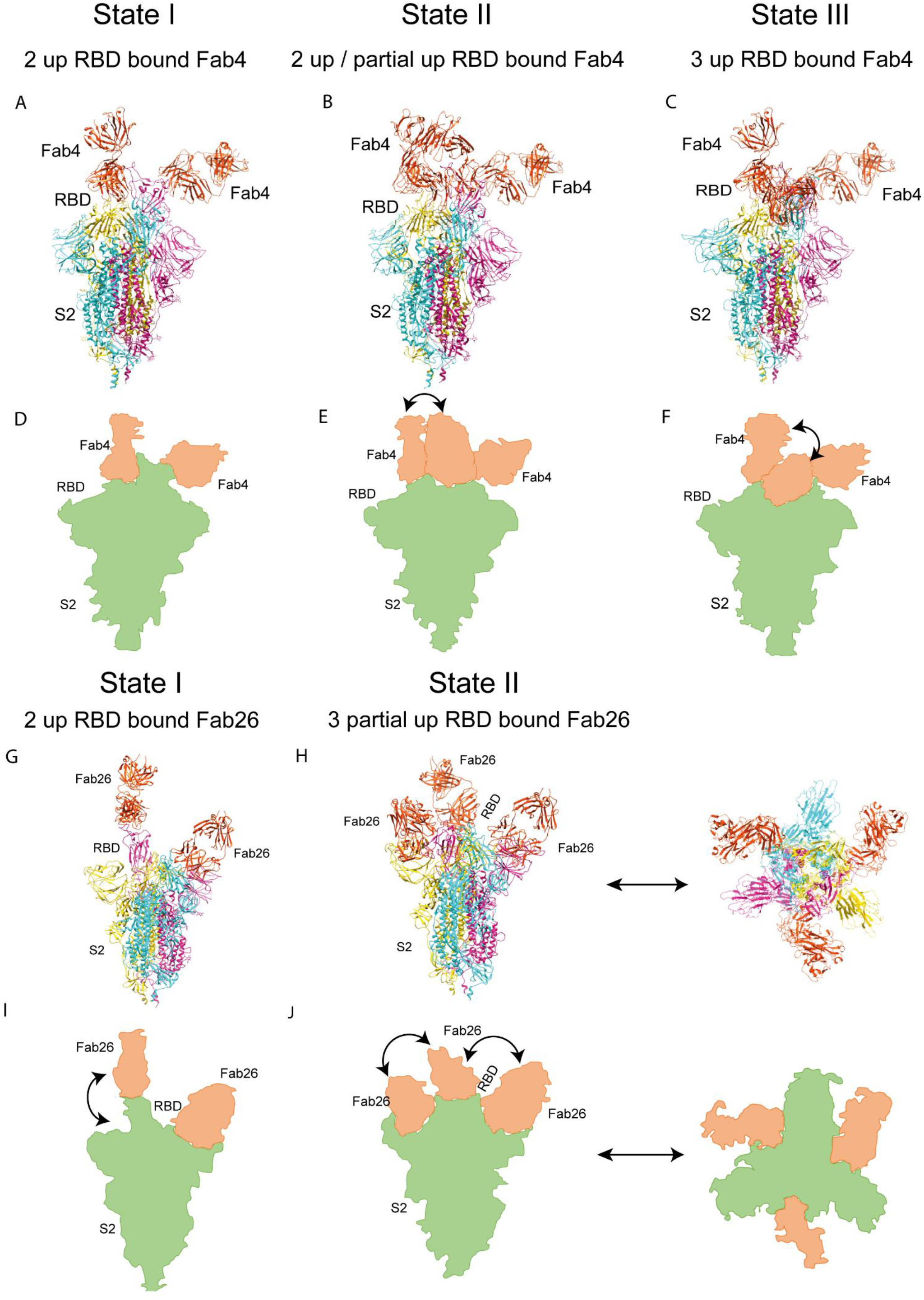
Stoichiometric analysis of different Fab4 (Left Panel) and Fab 26 (right panel) bound states: **A**. State I represent atomic model of two Fab4 bound to two “up” RBD conformation protomers. **B**. State II represents the atomic model of two Fab4 bound to “up” RBD and one Fab4 bound to partial open RBD form. **C**. State III represents atomic model of three Fab4 bound to open form of the protomers. Shifting of one RBD is shown in arrow while binding to the Fab4. **D, E, F**. Schematic diagram of both the states is shown to represent the RBD movement to accommodate three Fabs. **G**. State I represent atomic model two Fab26 bound to two “up” RBD conformation protomers. Shifting of one RBD is shown in arrow while binding to the Fab26. **H**. State II represents an atomic model of three Fab26 bound to partial open form of the protomers. In the extreme right, perpendicular views of state II are given. Shifting of all the RBD from each other shown in arrow upon binding to Fab26. **I, J**. Schematic diagram of both the states is shown to represent the RBD movement to accommodate three Fabs.

As stated above, among two states of Fab26 and S protein complexes, the 7.3 Å resolution state I showed two RBDs in up conformation bound to Fab and another RBD found in closed form, whereas the 4.8 Å resolution EM map of the state II exhibited three RBDs are interacting with Fab in the partial open conformation (Figure 2D, E; 5G-J). The crystal structure of open conformation of S protein (PDB ID: 6ZGG) was next fitted into the auto-sharpened EM map to generate atomic models for both states of the complex (Figure 4A; 5G-H). The CDR-RBD interfaces were observed to be similar across the state I and state II complex structures (Figure 5G-J). The common interacting residues found are A475, V483, E484, F486, and N487 (Figure 4B). To accommodate the third Fab26, a clear shift from “up” RBD to “partially open” was observed (Figure 5G-J). In the superimposed model of protomer 1 of both states, whereas the shift was minimal in the case of the second protomers of both the conformational states (Figure 5G-J). To measure the shift of interacting residues in both states, we selected one of the common interacting residues F486. A prominent 30 Å of F486 shift was observed in between the first promoter binding to two Fab26 and three Fab26. In the case of the second protomer, the shift is 43 Å, while the change of F486 position in the third protomer is 50 Å. These shifts suggest the stabilization of partial open conformation in state II to potentially neutralize the virus. The shift of protomer 1 and protomer 3 in the RBD region of state I as compared to state II was found to be 12.1° and 26.1° respectively, whereas no change was observed in between the RBD region of both the states. This inward movement of RBD in two protomers signifies the third protomer binding, facilitating partial open conformation-mediated binding of three Fab26s (Figure 5G-J). Cryo-EM-based structural studies show that RBDs are flexible, which could adapt the open conformation, as well as partial open conformation during the interaction with antibodies. The flexibility of RBD was supported by our previous studies (*15*). In addition, we have specified the interacting residues responsible for Fab-RBD stabilization, which also helped us to understand whether these two antibodies have affinities for other SARS-CoV-2 variants.

### Antibody binding alters the conformational states of SARS-CoV-2 S trimer in its apo and hACE-2 bound forms

To understand any conformational variations in the RBD region occurred after binding to Fab4 and Fab26 with respect to its apo form and in ACE2 bound complex, we compared our Fab4 and Fab26-S protein complex with hACE2 bound- and apo form of the S protomer. For this analysis, we used the first protomer of S trimer-Fab4 (Figure 3A) and-Fab26 (Figure 4A) high-resolution atomic model which was found to be most stable in all the conformations. In structural analysis, superimposition of both Fab’s bound complexes demonstrates its distal targeting to the RBD region independently (Figure 6E). Further, sequence analysis of potential epitope resides reveals the hotspot domain for each Fab’s with respect to different variants (Wuhan reference strain, wild, and omicron). The region of N439-V445 and V483-C488 residues forms a primary hotspot for antigen-antibody interaction for Fab4 and Fab26 respectively, which also validates the point that independent fashion antibody targeting (Figure 4C). In addition, structural analysis also proven its independent nature without any clashes (Figure 6, E). For Fab4, a 7.1° shift in the plane was observed as compared to the hACE2 bound protomer and an average of 8.2 Å shift in the ACE2 binding loop of the RBD spanning from N440 to V445 (Figure 6A, B). On the other hand, another loop containing two interacting residues P499 and T500 moved upwardly 9.6 Å to form contacts with Fab4. with respect to Fab26, a minor shift of 1.2° from the hACE2 bound structure and an average of 6.3 Å shift in the ACE2 binding RBD ridge were observed (Figure 6C, D). Q493 and Y489 have been reported to be pivotal residues from the RBD region in interaction with the ACE2 receptor (*35*). Interestingly, these two residues were found to be falling in the region of Fab26 interacting residues. Likewise, we observed 8.2 Å and 7.3 Å shifts in the respective residues Q493 and Y489, along with a 2.3° outward shift of the entire RBD core. The interfacial area of ACE2 and Fab4 is 3791 Å2, where 53 residues of heavy chain clashing with 46 residues of ACE2 containing 1505 area and 61 residues of light chain falling under 2286 interface area. Our analysis showed the overlapping interfacial area with ACE2 which emphasizes the inhibition of ACE2 to S trimer binding (Figure 6C, D).

**Fig 6:**
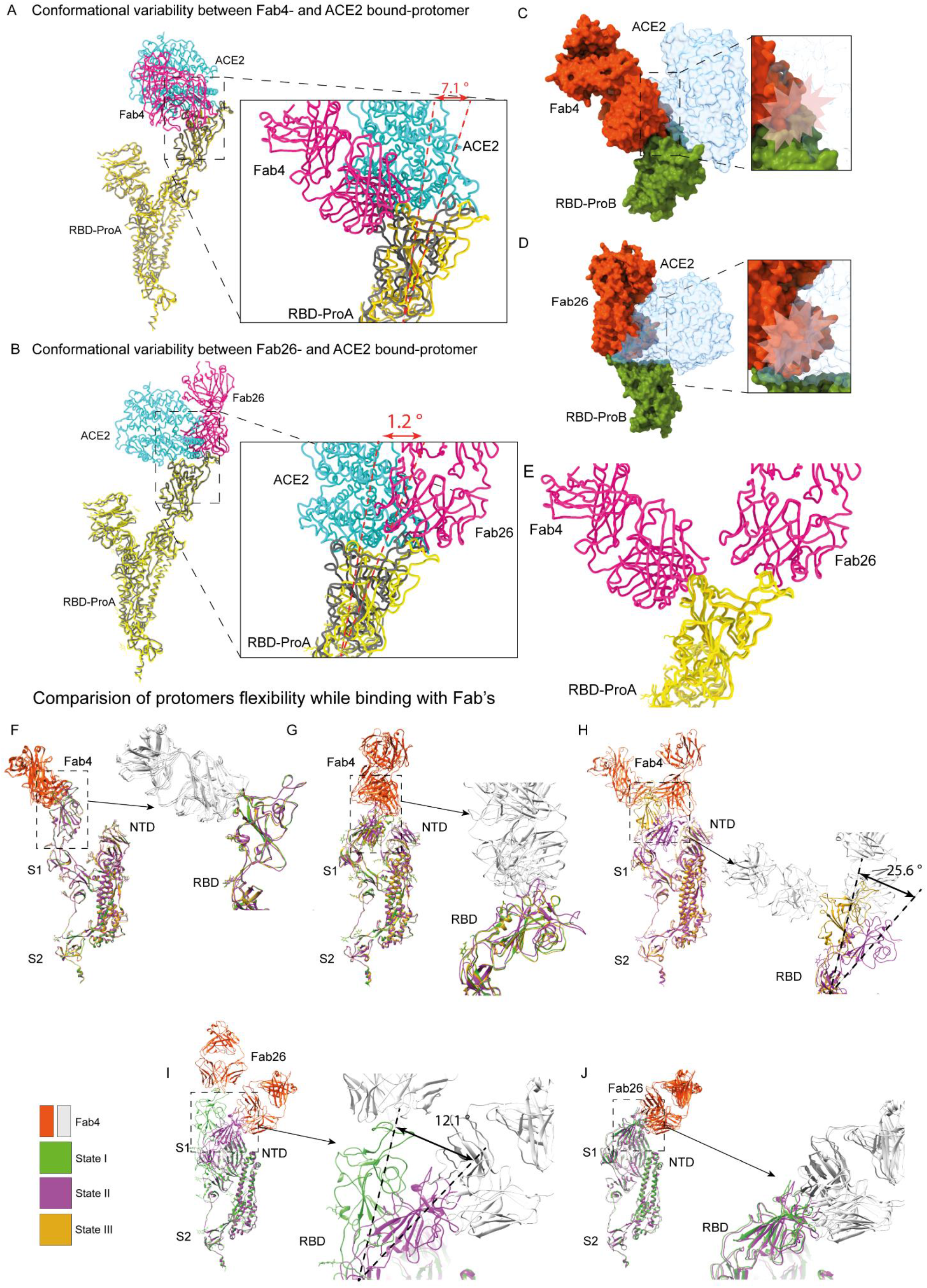
Comparison of conformational changes happens in S protein in Fab4 and Fab26 bound complexes: **A**. Licorice diagram of the Spike bound to Fab4 and ACE2 and zoomed view of conformational variable region of RBD bound to Fab4 and ACE2 is shown. ACE2 is shown in lime green, Fab4 in hot pink, black ribbon is Fab4 bound Spike and gold is ACE2 bound S protein. **B**. Licorice diagram of the conformationally variable region and the clashing region of Fab26- and ACE2 bound RBD. **C,D**. Clashing region of Fab and Fab26 bound RBD is shown in comparison with ACE2 bound RBD in silhouettes form. **E**. Distinctbinding loop for binding Fab4 and Fab26 in the RBD region is highlighted. **F-H**. Individual superimposition of protomer 1 (F), protomer 2 (G), and protomer 3 (H) from three different states of the Fab4-S protein complex displays the dynamics of each protomers across the states. Zoomed view of the RBD region is shown in every right corner of the enlarged view. **I-J**. Similarily, comparison of protomer 1 (I), and protomer 2 (J) in state I and state II of Fab26-S protein complex are demonstrated.

### Structure assisted modeling to predict correlates associated with resistance and sensitivity Omicron variants against THSC20.HVTR04 and THSC20.HVTR26

We finally examined the likelihood of the structural correlates associated with the neutralization resistance of Omicron BA.1 (B.1.1.529) against THSC20.HVTR04 Fab using our structural data. Towards this, superimposition of the S protein of Wu-1 (present study) and Omicron BA.1 (PDB ID: 7WBP) indicated conformational variations in the epitope of both Fab’s (Figure 7A, B). The primary interacting loop within the spike that forms epitopes for Fab4 (N439 to N450) and Fab26 (N481 to F490) were considered for the analysis. We observed a significant shift of 4.5° in the Fab4-Wuhan S protein interacting region in the context of the Omicron S protein and mild variations of 1.6 ° shift in the Fab26 binding domain. Further, MSA analysis of Wuhan reference strain wild (PDB ID: 6LZG), lately emergent significant variant strains globally: Delta (PDB ID: 7WBQ); Omicron BA. 1 (PDB ID: 7WBP) highlights the mutation in the epitope or nearby residues. Variations observed through structural and sequential analysis could be the possible cause for the evasion of omicron variant BA. 1. Biochemical studies found that Fab4 and Fab26 are having a null or moderate affinity with BA. 1 S protein. Interestingly, Fab4 shown moderate affinity with BA.2 omicron S protein, a dominating variant in India (Figure 7E). Sequence alignment of Wuhan RBD with Omicron variants BA.1, BA.2 and BA.4 RBDs and analysis of the atomic model of Fab4-WuRBD superimposed on crystal structure of BA.1, BA.2, and BA.4 (PDB ID: 7XBY, 7XO9, 7ZXU respectively) suggest that combined effect of N440K and G446S mutation causes BA.1 resistant to THSC20.HVTR04 (Figure 7A). While N440K mutation is common to all Omicron variants (BA.1, BA.2, and BA.4/BA.5), G446S mutation is present in BA.1 only and seems to be a vital residue to restrict THSC20.HVTR04 in neutralizing Omicron BA. 1. This observation was further supported by the fact that the presence of G446 in BA.2 and BA.4/BA.5 enabled THSC20.HVTR04 to efficiently neutralize them. Variations in electrostatic potential distributions along the surface of Fab’s interacting domains are highlighted across different strains (Figure 7C, D). Structural comparison using the atomic model of Fab26-wuRBD (Lime green) superimposed on the crystal structure of BA.1, BA.2, and BA.4 (PDB ID: 7XBY, 7XO9, 7ZXU respectively) (magenta color) suggest that T478K mutation likely abrogating the ability of THSC20.HVTR26 to bind to BA.2 (Figure 7B). Finally, structural and neutralization data suggest that the appearance of F486V mutation has a profound effect on less surface charge localization at the epitope site of RBD associated with the inability of binding of THSC20.HVTR26 Fab to BA.4 spike expressed on cell surface (Figure 7B, D). Besides, some key point mutations in the epitope region change in surface charge distribution also contribute for the decreased affinity of both mAbs with omicron (Figure 7C, D). A combination of all the above features would likely contribute in conferring resistance of Omicron variants to these two novel mAbs.

**Fig 7:**
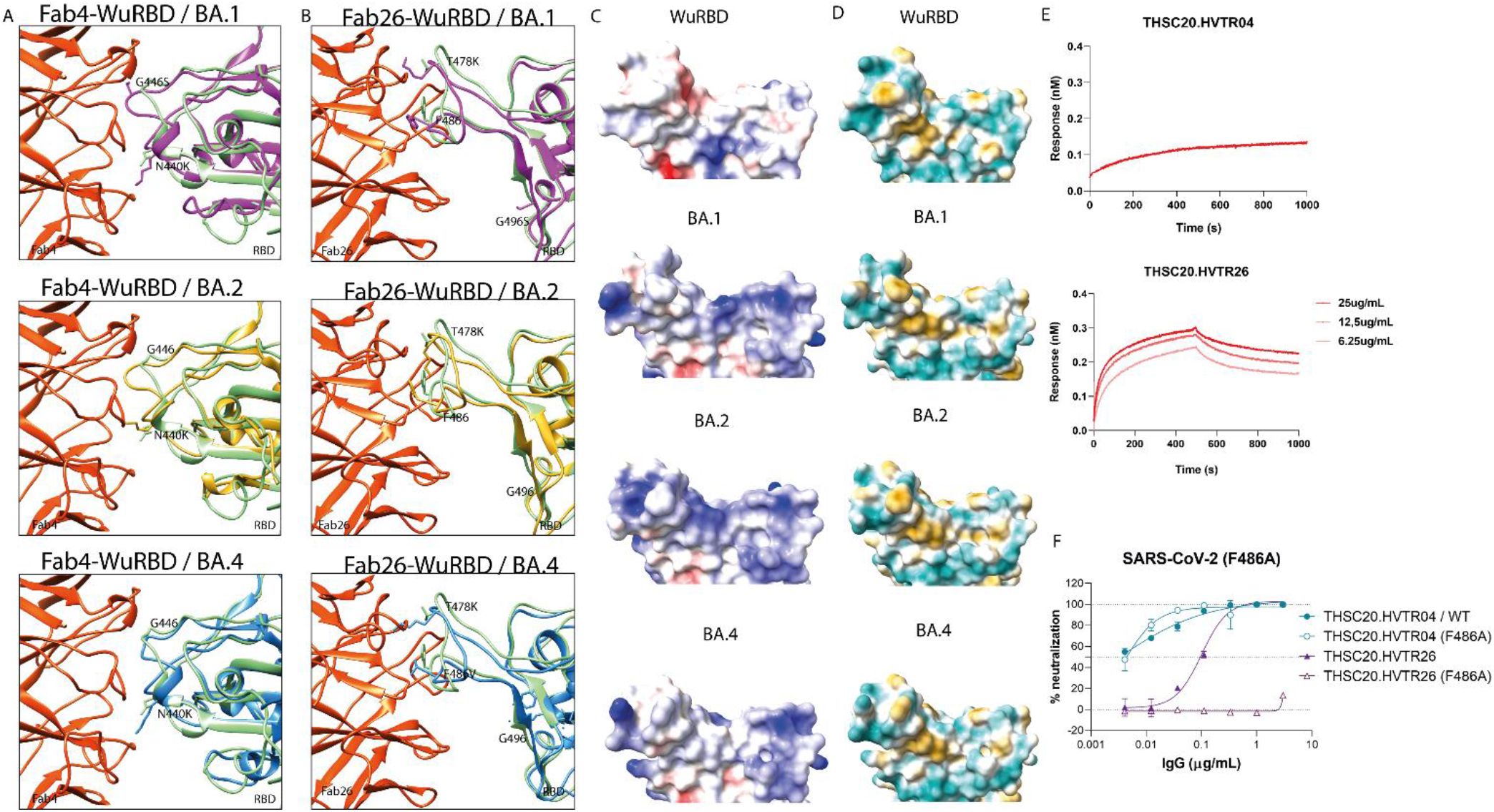
Mutation-driven alteration in the epitope region associated with BA.2, BA.4/BA.5 (but not BA.1) neutralization by mAb4 and mAb26. **A**. Superimposition of wild-type RBD (lime green) bound to mAb4 (orange red) and RBD (magenta) of BA.1 strain shows changes in two crucial interacting residues G446S, N440K results in weak interaction of mAb4 with BA. 1 spike protein. Whereas, superimposition of wild type RBS with BA.2 (gold) and BA.4 RBD (dodger blue) shows due to changes in only one interacting residues mAb4 has moderate affinity towards BA.2 and BA.4. **B**. Superimposition of wild-type RBD (lime green) bound to mAb26 (orange red) and RBD (magenta) of BA. 1 strain shows changes in two crucial interacting residues T478K, G496S resulting in weak interaction of mAb26 with BA.1 spike protein. Similarly, changes in the residues of BA.2 RBD (gold) T478K and BA.4 RBD (dodger blue) T478K and F486V, strikingly altered the affinity of mAb26 towards newer circulating variants. **C-D**. Electrostatic surface potential map and hydrophobicity mapping of the RBD region of different SARS-CoV 2 variants including the latest circulating variants suggest the changes in the surface charge due to some key point mutations results in the differential affinity pattern of mAb4 and mAb26 towards the RBD of these variants. **E**. Binding studies of both the mAbs against Omicron BA.1 spike using BLI-octet. **F**. Effect of F486A mutation in the Spike of SARS-CoV-2 on the neutralization potential of mAbs THSC20.HVTR04 and THSC20.HVTR26.

## Discussion

The outbreak of SARS-CoV-2 and its continued emergence of mutated strains became the global thread for the recent years. In particular, its impact on the function and neutralization efficiency of antibodies. The majority of potent monoclonal antibodies reported till date against SARS-CoV-2 lost their neutralizing efficiency for emerging variants (VOC) with the exception of few antibodies (*36, 37*). In that order, two promising antibodies THSC20.HVTR04 (Fab4) and THSC20.HVTR26 (Fab26) screened from convalescent sera of wild type infected patient shown to be highly effective against majority of SARS-CoV-2 VOC’s. Though the latest emerged variant Omicron displays a high mutational and transmission rate, several studies reported its case fatality rates (CFR) and need for hospitalization, etc. is much lower than seen in the previous variants. Interestingly, the antibody THSC20.HVTR26 (Fab26) shows moderate neutralization against the Omicron BA. 1 variant. Cryo-EM study of S protein with two potent antibodies provides a platform to understand the dynamic flexibility of Fab’s interaction network and the precise epitopes. The heavy chain of both Fab’s predominantly interacts with Spike RBD. In this work, we are reporting the structural accommodation and mechanism of two potent SARS-CoV-2 antibodies in complex with the native form of the ancestral SARS-CoV-2 spike protein. The quantification of epitope-targeting frequency were analyzed between the different conformations and deduced the potential epitopes for both the Fab’s. It helps in understanding the mechanism of viral escape from neutralizing antibodies, particularly for the recent variant omicron. Furthermore, analysis of RBD sequences of different variants provides an insight into the amino acid modifications in the binding domain of the mAbs studied here. The key epitopes N440K (Fab4) and E484A (Fab26) were found in the modified residues with respect to the variant Omicron. Likewise, we observed a divergence in the surface map of electrostatic and hydrophobicity over the paratopes of Wuhan and Omicron variants BA.1, BA.2, BA.4. It is likely that a combination of these factors is likely to be the reason for the reduction of neutralization efficiency against Omicron variants.

The proposed hypothesis of S protein binding to Fab4 are; a. In solution, the spike trimers exist in the equilibrium between closed and partially open conformations (*14*, *15*). Initially, two Fabs bind to protomers I and II (state I) (Figure 5A, D). Following, another Fab4 bind to the third protomer in the partial open conformation (Figure 5B, E). This stage behaves like an intermediate stage where a clash happens between the third Fab4 to the second protomer, resulting in a slight change of conformation in the Fab4-protomer 2 complex. Later on, Fab4 on the third protomer is likely to be stabilized at upright conformation at stage III (Figure 5C, F). Acquiring protomer 3 in upright conformation is likely to prevent the Fab and protomer 2 happens in state II. The initial state I (Figure 5A, C) and final state III (figure 5D, F) seem to be very less deviation and signify possibly a stabilized state. In a similar context, in the Fab26-S protein complex the partial open conformations of are stabilized in the three Fab26 bound states with the S trimer. At first, two Fab26 are attached to the epitope region of the RBD in “up” conformation. To facilitate the binding of the third Fab26, protomer 1 and protomer 3 are shifted from “up” form to partial open form. Therefore, the deviation in the RBD loop to adopt a thermodynamically stable partial open form is observed to accommodate the third Fab26 in the protomer 3 positions. In summary, through atomic-level structural characterization via high resolution cryo-EM, our study highlighted key features of interactions of the two novel mAbs with the SARS-CoV-2 spike protein that distinguishes them in their ability to cope up in tackling the Omicron lineage variants and which opens up the possibility of structure guided modification of these mAbs towards overcoming the evolving mutations in the viral spike protein.

## Materials and Methods

### Preparation of IgG and Fabs

IgGs were prepared following the reported protocols (*33*). The Fragment Antigen Binding (Fabs) of each IgG were prepared by using ~ 500 μl of antibody (mAb) (1-5 mg/ml) was mixed with 1200 μl of 100mM Sodium Acetate (sodium acetate anhydrous from Sigma Aldrich) having 1mM Ethylenediaminetetraacetic acid disodium salt dihydrate (EDTA for electrophoresis, for molecular biology, 99.0-101.0% from Sigma), 100 μl of 50 mM Cysteine (L-Cysteine for biochemistry from Merck) and 200 μl of (in 10 μg/mg mAb concentration) Papain (Papain from papaya Latex, 89% pure lyophilized powder from Sigma). The mixture was then kept in the incubator shaker at 37 degrees Celsius for 8-10 hours. After 10 hours 75mM (~150 μl) Iodoacetamide (Iodoacetamide BioUltra from Sigma) was added to the mixture to stop the papain activity and incubated at room temperature for 30-40 mins. Then the sample was added to the protein A beads (Immobilized Protein A resin from G Biosciences) washed 2-3 times with PBS to remove the storing ethanol and incubated for another 3-4 hours at room temperature on the rocker. After ~ 4hrs the mixture was poured into the gravity column and the flow through was collected in a 15 ml falcon. The flow through contained the Fab region whereas FC region was bound to the beads. The beads were then washed with PBS twice to collect any Fab fractions out from the column. The column was then eluted with 100 mM glycine (G Biosciences), pH=2.2 to remove the bound Fc fractions and then was washed with PBS for 3-4 times. Finally, the flow through and the wash PBS containing any residual Fab fractions were pooled together and brought to a final concentration of 0.5-0.8 mg/ml.

### Purification of SARS-CoV-2 RBD

His-tagged SARS-CoV2-RBD plasmid DNA construct was kindly provided by Dr Jason S. McLellan’s (*41*). The His tagged RBD construct was purified from the culture supernatant of Expi293 cells (Thermo Fisher) that were transfected with the plasmid DNA using FectoPRO (Polyplus Inc.). Culture supernatants were harvested between 4-5 days and the RBD protein was purified from filtered cell supernatants using Ni-NTA resin (G Biosciences, Inc.) following the protocol as has been described by McLellan *et al*. (*41*) The Ni-NTA column was equilibrated with PBS and washed Ni-NTA bound RBD with 10mM imidazole (Sigma Aldrich) containing PBS (pH 7.4). The RBD protein was finally eluted using 300 mM imidazole. The eluted protein was buffer exchanged with PBS (pH 7.4) and was concentrated using 10 kDa cut off Amicon (Merck Millipore) concentrator to a final concentration of 1 mg/ml and stored at −80°C until further use.

### Spike 6P protein Expression and Purification

SARS-CoV-2 S HexaPro plasmid was a gift from Jason McLellan (Addgene plasmid # 154754). This plasmid contains CMF promotor driven expression of the SARS-COV-2 Spike-B.1 ectodomain (1-1208 AAs) with mutated furin site (682-685 GSAS) and Hexa proline mutations F817P, A892P, A899P, A942P, K986P, V987P as foldon. The HRV 3C cleavage site was placed before tags 8X His tag followed by 2X Strep-Tag II at the C terminal. (*42*)

Plasmid was transfected to suspension Expi293F cells using Expifectamine 293 Transfection Kit (Gibco, Thermo Fisher, Cat # A14524) according to the manufacturer’s instruction. In brief, 24 h prior to transfection, 100% viable Expi293F cells were passaged at a density of 2 million per ml. On next day 4 million per ml cells (100 ml) were transiently transfected with Expifectamine-plasmids-DNA complex (100 μg plasmid, complexed with 270 μl of ExpiFectamine293). Post 20 h, Enhancer 1 and Enhancer 2 were added according to the manufacturer’s protocol. After five days from day of transfection, culture supernatant without cell was collected and used for purification of 6X Histidine tagged protein. Media supernatant were affinity purified by immobilized metal affinity chromatography using Ni-NTA resin (G Biosciences, Cat # 786940). 100 ml media supernatant was mixed with PBS (pH 7.4) equilibrated Ni-NTA resin (5 ml) and incubated for 4 h at 4°C for protein immobilization under gentle rotation condition. The Supernatant resin complex was gently applied to column. After unbound fraction collection, A ten-column volume wash of 1× PBS (pH 7.4), supplemented with 25 mM imidazole was performed. Finally, the bound protein was eluted with a gradient of 200 to 500 mM imidazole in PBS (pH 7.4). The eluted fractions were pooled and dialyzed against 1X PBS buffer (pH 7.4) twice using dialysis tubing cellulose membrane [avg. flat width 43 mm (1.7 in.), 14kDa MWCO, Sigma-Aldrich, Cat # D9527]. Protein concentration was determined by absorbance (A280) using Biophotometer D30 (Eppendorf) with the theoretical molar extinction coefficient (146720 M-1 cm-1) calculated using the ProtParam tool (ExPASy). The protein was analyzed on 8% SDS PAGE for purity and homogeneity under reducing and non-reducing conditions.

### Site-directed mutagenesis

Point substitutions within RBD in SARS-CoV-2 spike gene were introduced by site-directed mutagenesis using the QuikChange II kit (Agilent Technologies Inc.) following the manufacturer’s protocol and by overlapping PCR strategy as described previously (*33*). Successful incorporation of desired substitutions was confirmed by Sanger sequencing.

### Cell surface binding assay

The binding of mAbs to the SARS-CoV-2 spikes expressed on the HEK 293T cell-surface was assessed as described previously with some modifications (*43*). Briefly, HEK293T cells were transfected with the three plasmids used to generate SARS-CoV-2 pseudovirus (SARS-CoV-2 MLV-gag/pol, MLV-CMV-luciferase and SARS-CoV-2 spike plasmids). After incubation for 36-48 h at 37°C, cells were trypsinized and a single cell suspension was prepared which was distributed into 96-well U bottom plates. 3-fold serial dilutions of mAbs starting at 10 μg/ml and up to 0.041 μg/mL were prepared in 50 μl/well and added to the spike expressing as well as un-transfected 293T cells for 1 hour on ice. Cells were subsequently washed twice with FACS buffer (1x PBS, 2% FBS, 1 mM EDTA) and then stained with 50 μl/well of 1:200 dilution of R-Phycoerythrin AffiniPure F(ab’)_2_ Fragment Goat Anti-Human IgG, F(ab’)_2_ fragment specific antibody (Jackson ImmunoResearch Inc.) for 45 min. Cells were finally stained firstly with 1 LIVE/DEAD fixable aqua dead cell stain (ThermoFisher) in the same buffer for another 15 minutes and subsequently washed twice in plates with FACS buffer. The binding of mAbs to spikes expressing on cell surface was analyzed using flow cytometry (BD Canto Analyzer). Percent (%) PE-positive cells for antigen binding were calculated and the binding data were generated. CC12.1 (SARS-CoV-2 mAb), and CAP256.VRC26.25 antibody (HIV-1 bnAb) were used as positive and negative controls respectively for this experiment.

### Biolayer Interferometry (BLI)

The binding affinity of the Fabs were assessed by using Biolayer interferometry (BLI-Octet). Ni-NTA biosensors (Forte’ Bio) were used to assess the binding affinities of Fab4 and 26 with SARS-CoV-2 RBD in PBST (PBS containing 0.02% Tween 20) at 30°C and 1,000 rpm. shaking on an Octet RED 98 instrument (Forte’ Bio Inc.). Sensors were first soaked in PBS for 15 minutes before being used to capture His-tagged SARS-CoV-2 RBD protein. RBD was loaded to the biosensors up to a level of 1.0 nm. Biosensors were then immersed into PBST for 100 seconds and then immersed into wells containing different concentrations of a Fab dissolved in PBST (PBS containing 0.02% Tween 20) for 500 seconds to measure association. A threefold dilution series with five different concentrations (600, 200, 66.7, 22.3, and 7.4 nM) was prepared for each Fab. Biosensors were next dipped into wells containing PBST for 500 seconds to measure dissociation. Data were reference-subtracted and aligned to each other using Octet Data Analysis software v10 (Forte’ Bio Inc.) based on a baseline measurement. Curve fitting was performed using a 1:1 binding model and data for all the five concentrations of Fabs. Kon, Koff and *K*D values were determined with a global fit.

### DLS and DSC

The particle size and thermal stability of the purified IgGs were characterized by Dynamic Light Scattering (DLS) and Differential scanning calorimetry (DSC) respectively. DLS was done using ~200 μl of 1mg/ml solution of the antibody in Nano-ZS90 instrument from Malvern and DSC was done using 750 μl of 1mg/ml concentration of protein in NANO DSC instrument from TA.

### Size exclusion chromatography (SEC)

The tendency of aggregation of the IgGs representing all the mAbs were characterized by SEC. For SEC), the samples were passed through Superdex™ 200 Increase 10/300 GL size exclusion column (GE, Inc.) and were eluted with degassed PBS buffer (pH 7.4) at 0.3 ml/min flow rate using AKTApurifier™ 100 (GE, Inc.).

### Sample preparation for Negative staining TEM and data processing

To analyze the binding of Fab to S protein and its homogeneity, we performed conventional negative staining TEM. The diluted samples of S protein (1 mg/ml) - 10 times and the Fab’s (1.4 mg/ml) - 300 times were mixed and incubated for 2 minutes. The 3.5 μl of sample mixture were applied onto freshly glow discharged carbon coated Cu grids (30 secs) (EM grid, 300 mesh, Electron Microscopy Sciences). After 1 min, the excess solvent was blotted and 1% uranyl acetate (Uranyl Acetate 98%, ACS Reagent, Polysciences, Inc.) was added on the grid. After air dried, the grids were used for data acquisition with room temperature 120 kV Talos L120C electron microscope. Data collection was performed using 4k x 4k Ceta camera at the magnification of 73kx and its calibrated at 3.84 Å/pixel. The collected micrographs were processed in EMAN 2.1 (*44*). Automated and manual mode of particle picking was performed, and its co-ordinates were extracted using e2boxer.py in EMAN 2.1. Subsequently, reference free 2D class averages allowed us to visualize various projections Fab’s bound S protein complexes. The cleaned dataset of the samples was used for reference-free 2D classification, and reference-free 2D class averages of different particle projections were calculated using simple_prime2D of SIMPLE 2.1 software (*45*) with a mask diameter of 30 pixels at 3.84 Å/pix.

### Cryo-EM Sample Preparation

R1.2/1.3 300 mesh gold grids (Quantifoil) (Electron Microscopy Sciences) were glow discharged for 90 seconds at 20mA before freezing. Equal volume of SARS-CoV 2 S protein (1 mg/ml) was mixed with 30 X diluted Fab’s (1.4 mg/ml) and incubated at room temperature for 2 minutes. Three microliters of final sample were applied onto the freshly glow discharged grids, incubated for 10 secs and immediately blotted for 8.5 secs at blot force of zero in pre-equilibrated chamber of FEI Vitrobot Mark IV plunger. Immediately after blotting, grid was plunged into the liquid ethane.

### Cryo-EM Data Collection

Cryo-EM data were acquired using 200 kV Talos Arctica transmission electron microscope (Thermo Scientific™) equipped with Gatan K2 Summit Direct Electron Detector. Movies were recorded automatically using Latitude-S (DigitalMicrograph - GMS 3.5) at nominal magnification of 54,000x at the effective pixel size of 0.92 Å (*46*). Micrographs are collected in counting mode with a total dose of 50 e^-^ /Å2, with an exposure time of 8 sec distributed for 20 frames. A total of 3789 and 2102 movies were acquired for the Fab4- and Fab26-S protein complexes respectively.

### Cryo-EM Data Processing

Single-Particle Analysis (SPA) were performed for the acquired cryo-EM data using the Relion version 3.1 (*47*). In the beginning, drift and gain corrections of the individual movies were corrected with MotionCor2 (*48*). Subsequently, motion corrected micrographs were subjected to screen for bad micrographs using cisTEM (*49*) and the detected fit resolution threshold chosen for screening is 7 Å. The best micrographs were chosen for an estimation of Contrast transfer function (CTF) parameters using CTFFIND 4.1.13 (*50*). Followed by, particles were picked by Laplacian picker in Relion and extracted with the box sizes of 336- and 360 Å for the S:Fab4 and S:Fab26 complexes, respectively. The well-defined classes for S:Fab4 and S:Fab26 complexes were obtained as 4,18,672 and 2,55,039 particles respectively after six rounds of 2D classification. These particles were selected for 3D classification without imposing symmetry (C1). To achieve high resolution, all particles belonging to best classes of Fab complexes were subjected to movie refinement, which includes estimation of beam tilt, anisotropic magnification and per-particle CTF refinement for defocus and astigmatism. The sharpening for the 3D auto-refined maps was performed with Relion 3.1 (*47*) and PHENIX (*51*, *52*). Overviews of cryo-EM data processing is shown in Figure S4, S5. Global resolution of Fourier shell correlation (FSC) were estimated at the threshold of 0.143 and the estimation of local resolution were performed with ResMap, using auto-refined half maps.

### Model building and structure refinement

Automated model building was iteratively performed with Phenix Real Space Refinement. Based on the S protein conformations, the SARS-S protein PDB: 7kms – 3 RBD up; 6zwv – 3 RBD down; 7akd-1 partial open RBD were docked with cryo-EM maps using UCSF Chimera “Fit in map” tool. To build the model for Fab’s, the query sequences of both chains were submitted to Swiss-Model and the resultant model were also docked to the EM maps. The fitted models were used as initial models and refined against sharpened EM maps. The structural statistics for Cryo-EM map and atomic model were analyzed using Phenix, EMringer, Molprobity, and UCSF chimera (*53*).

### Analysis and Visualization

Cryo-EM maps and atomic models were visualized using UCSF Chimera. PDBsum was used to identify the interacting residues of S protein and Fab complexes (*54*). Surface coloring based on Kyte-Doolittle hydrophobicity scale were performed using UCSF ChimeraX (*55*). Angle of chain rotation were estimated by creating the planes between three terminal residues (N370, F490, P1140) of spike protomer and its shift is calculated between the planes. The RMSD values were calculated using the UCSF Chimera “MatchMaker” tool.

## Acknowledgments

We thank Mynvax Private Limited for support with SARS-CoV-2 Spike protein expression and characterization. We acknowledge the Department of Biotechnology, Department of Science and Technology (DST) and Science, and Ministry of Human Resource Development, India for funding the cryo-EM facility at IISc-Bangalore. We acknowledge DBT-BUILDER Program (BT/INF/22/SP22844/2017) and DST-FIST (SR/FST/LSII-039/2015) for the National Cryo-EM facility at IISc, Bangalore. We acknowledge the financial support from the Science and Engineering Research Board (SERB-EMR/2016/000608, and SERB-IPA/2020/000094) for financial support for consumables, and DBT-IISc partnership program phase II for the negative staining TEM facility at Biological Sciences Division, IISc. We thank Penny Moore and Jinal Bhiman, NICD, Johannesburg, South Africa to kindly provide us with the Omicron spike plasmid constructs, Jason McLellan, Texas, USA for kindly providing us with the SARS-CoV-2 RBD construct used for purification of RBD protein and Devin Sok for valuable suggestions and inputs. This study was supported by funding from the Research Council of Norway (Project ID: 285136), Bill and Melinda Gates Foundation supported Global Immunology & Immune Sequencing for Epidemic Responses (GIISER-South Asia) project at THSTI, (INV-030592). We thank Prof Pramod K Garg, Executive Director, THSTI and lead PI of the BMGF-GIISER-South Asia project for support. JB is supported by the DBT-Wellcome Trust India Alliance Team Science Grant (IA/TSG/19/1/600019).

## Declaration of interest

A patent application (PCT/IB2022/057923) filed on the invention of novel monoclonal antibodies with the following inventors: Jayanta Bhattacharya, Nitin Hingankar, Payel Das, Suprit Deshpande, Pallavi Kshetrapal, Ramachandran Thiruvengadam, Amit Awasthi, Zaigham Abbas Rizvi

## Authors contribution

SD designed the TEM and cryo-EM structural studies. CFR and AC performed the negative staining sample preparation and image processing. CFR and AC carried out cryo-EM samples preparation, data acquisition, data processing and structural analysis. MYA and SM prepared the antibody Fabs, purified RBD, carried out BLI assay and characterized the biophysical and biochemical properties of the mAbs. NH and PD helped in cloning and purification of mAbs; SuD and DR carried out cell surface binding of mAbs with SARS-CoV-2 spikes expressed on cell surface by FACS; SuD carried out site-directed mutagenesis and pseudovirus neutralization assay.

## Deposited Data Accessibility

Cryo-EM density maps of Wuhan SARS-CoV-2 trimeric spike protein complexed with monoclonal antibodies (THSC20.HVTR04 & THSC20.HVTR26) were deposited in Electron Microscopy Data Bank (EMDB). The various conformational states captured for each antibodies and its identifiers at EMDB are:

|  | EMDB accession code(s) |
| --- | --- |
| Spike- THSC20.HVTR04(Fab4) complexes |  |
| State - I: Spike 2-up RBD with THSC20.HVTR04 (Fab4) | EMD-34546 |
| State - II: Spike 3-up RBD with THSC20.HVTR04 (Fab4) | EMD-34547 |
| State - III: Spike 3-up RBD with THSC20.HVTR04 (Fab4) | EMD-34548 |
| Spike- THSC20.HVTR26(Fab26) complexes |  |
| State - I: Spike 2-up RBD with THSC20.HVTR26 (Fab26) | EMD-34563 |
| State - I: Spike 2-up RBD with THSC20.HVTR26 (Fab26) | EMD-34602 |
| State - II: Spike 3-up RBD with THSC20.HVTR26 (Fab26) | EMD-34603 |

## Notes

### Competing Interest Statement

The authors have declared no competing interest.

