## Supplementary File for "High resolution cryo-EM structures of two potently SARS-CoV-2 neutralizing monoclonal antibodies of same donor origin that vary in neutralizing Omicron variants"

**Supplemental Information**

**Table S1**: Cryo-EM Data collection, image processing and refinement for S Protein: Fab4 complex

| **Data Collection and Processing** | **State I** | **State II** | **State III** |
|---|---|---|---|
| Magnification | 54,000 kX | 54,000 kX | 54,000 kX |
| Voltage | 200 kV | 200 kV | 200 kV |
| Electron exposure ($e^-$/Å$^2$) | 50 | 50 | 50 |
| Defocus range (µm) | -1.25 to -2.75 | -1.25 to -2.75 | -1.25 to -2.75 |
| Pixel size (Å) | 0.92 | 0.92 | 0.92 |
| Symmetry Imposed | C1 | C1 | C1 |
| Number of Movies | 3789 | | |
| Number of Particles | 1,63,492 | 93,959 | 49,160 |
| Map Resolution (Å) | 4.54 | 5.152 | 4.9 |
| FSC threshold | 0.143 | 0.143 | 0.143 |
| Map Resolution Range | 3.3-6.1 | 3.3-6.1 | 3.3-6.1 |
| Map Sharpening B Factor (Å$^2$) | -273 | | |
| RBD Conformation | 2-RBD Up | 3-RBD Up | 3-RBD Up |

**Table S2:** Cryo-EM Data collection, image processing and refinement for S Protein: Fab26 Complex

| **Data Collection and Processing** | **Single Fab Masked** | **State I** | **State II** |
|---|---|---|---|
| Magnification | 54,000 kX | 54,000 kX | 54,000 kX |
| Voltage | 200 kV | 200 kV | 200 kV |
| Electron exposure ($e^-$/Å$^2$) | 45 | 45 | 45 |
| Defocus range (µm) | -1.25 to -2.75 | -1.25 to -2.75 | -1.25 to -2.75 |
| Pixel size (Å) | 0.92 | 0.92 | 0.92 |
| Symmetry Imposed | C1 | C1 | C1 |
| Number of Movies | 2102 | | |
| Number of Particles | 1,09,596 | 26,719 | 48,584 |
| Map Resolution (Å) | 4.4 | 7.3 | 4.8 |
| FSC threshold | 0.143 | 0.143 | 0.143 |

| | | | |
|---|---|---|---|
| Map Resolution Range | 3.3-6.1 | 3.3-6.1 | 3.3-6.1 |
| Map Sharpening B Factor (Å$^2$) | -249 | | |
| RBD Conformation | 2-RBD Up | 2-RBD Up | 3-RBD Up |

MolProbity Score for S Protein: Fab4 Complex

| | SARS-CoV-2 + Fab4 | | |
|---|---|---|---|
| **Validation** | **State I** | **State II** | **State III** |
| MolProbity Score | 1.88 | 2.28 | 2.32 |
| Clash Score | 7.36 | 18.31 | 18.64 |
| Rotamer Outliers (%) | 0.20 | 0.26 | 0.18 |
| Ramachandran Plot | | | |
| Outlier (%) | 0.11 | 0.16 | 0.18 |
| Allowed (%) | 7.66 | 8.73 | 9.78 |
| Favored (%) | 92.23 | 91.11 | 90.04 |

MolProbity Score for S Protein:Fab26 Complex

| | SARS-CoV-2 + Fab26 | | |
|---|---|---|---|
| **Validation** | **Single Fab Masked** | **State I** | **State II** |
| MolProbity Score | 1.99 | 3.03 | 2.11 |
| Clash Score | 8.77 | 126.30 | 11.39 |
| Rotamer Outliers (%) | 0.40 | 0.09 | 0.54 |
| Ramachandran Plot | | | |
| Outlier (%) | 0.14 | 0.10 | 0.07 |
| Allowed (%) | 8.98 | 7.30 | 9.43 |
| Favored (%) | 90.88 | 92.60 | 90.51 |

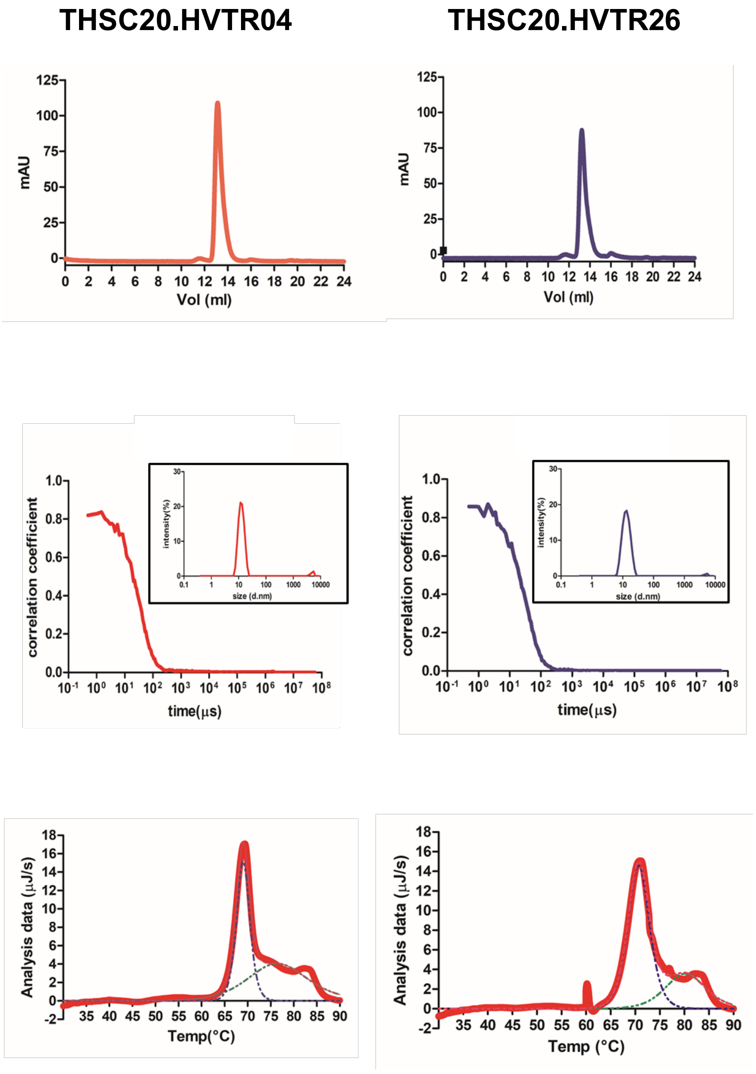


**Fig S1.** Biophysical characterization of mAb4 and mAb26 for its purity and stability. A. Size exclusion chromatography studies of mAb4 and mAb26. The mAbs eluted at ~ 13.5 ml respectively as a monomer. The data suggest that the purified mAbs are pure and devoid of any aggregates and other high or low molecular weight species. B. DLS (dynamic light scattering) studies of mAb4 and mAb26. The correlation coefficient function show single decay component. The size distribution by intensity (inset) shows a single effective peak of diameter around 12.93nm and 13.57nm respectively, with negligible evidence of aggregates or other components. The data suggest that the purified mAbs are homogeneous. C. DSC (differential scanning calorimetry) studies of mAb4 and mAb26. The DSC thermogram of mAb4 and mAb26 with majorly Fab domain estimated the melting temperature of Fab domain of mAb4 and mAb26 to be 69.03ºC and 70.74ºC respectively. The data suggest that the mAbs are thermodynamically stable.

Supplementary figure 2

A.

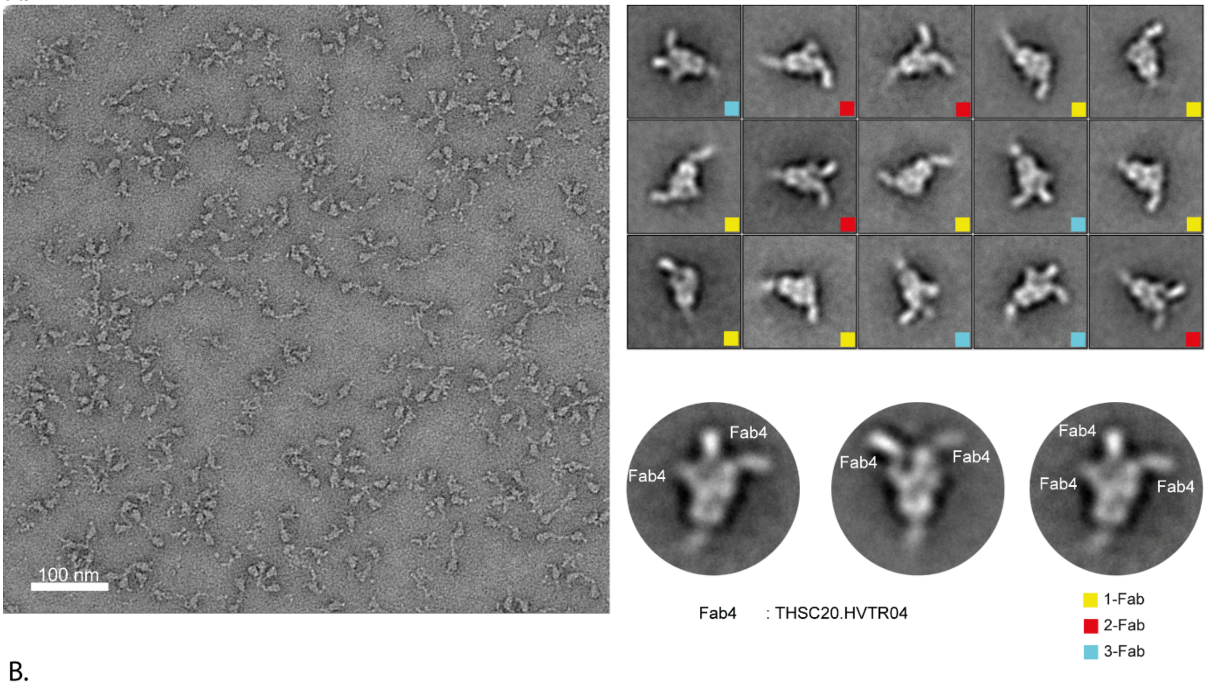


B.

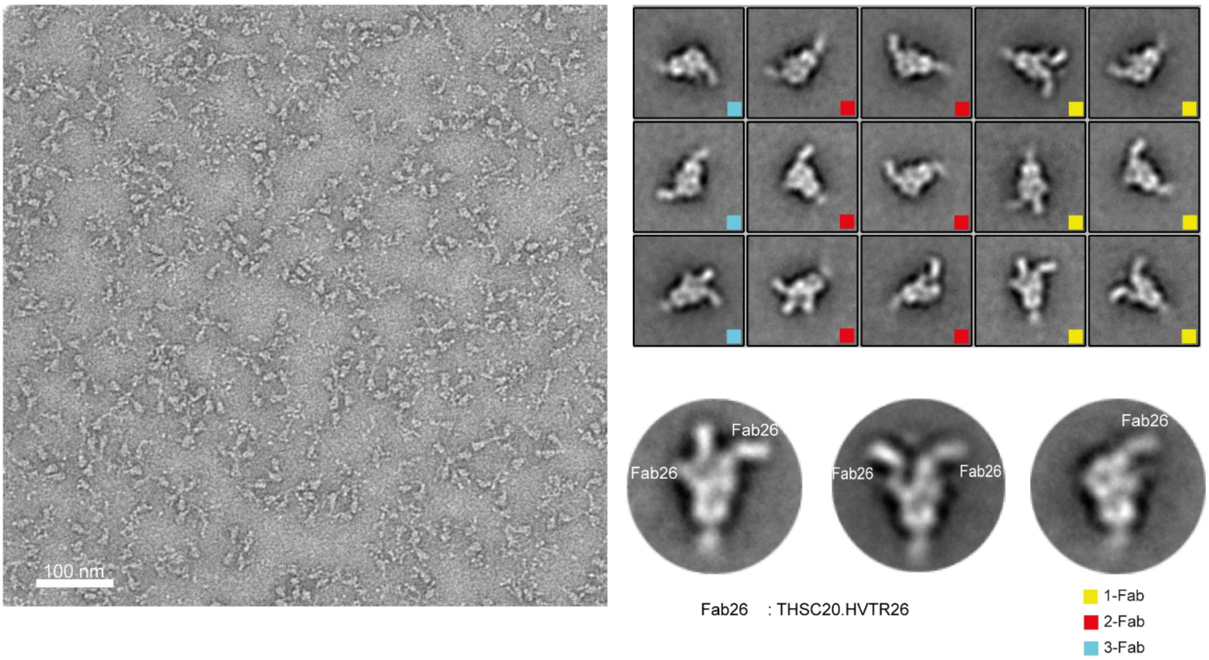


Figure S2: Negative staining micrograph and 2D class averages of S protein with A. Fab4 and B. Fab26 complexes

Supplementary figure 3

A.

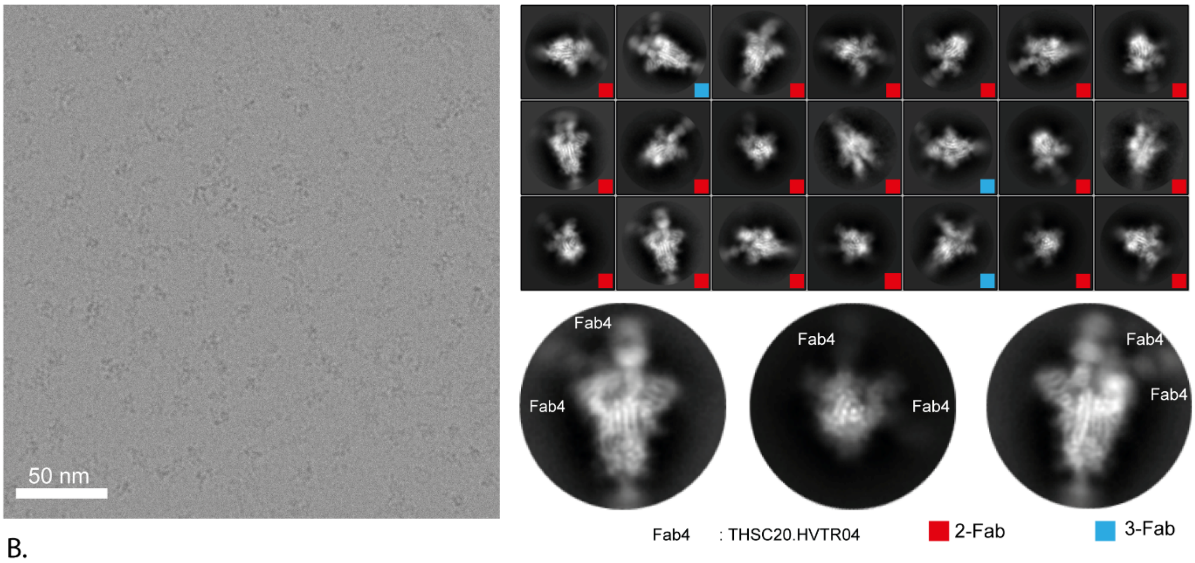


B.

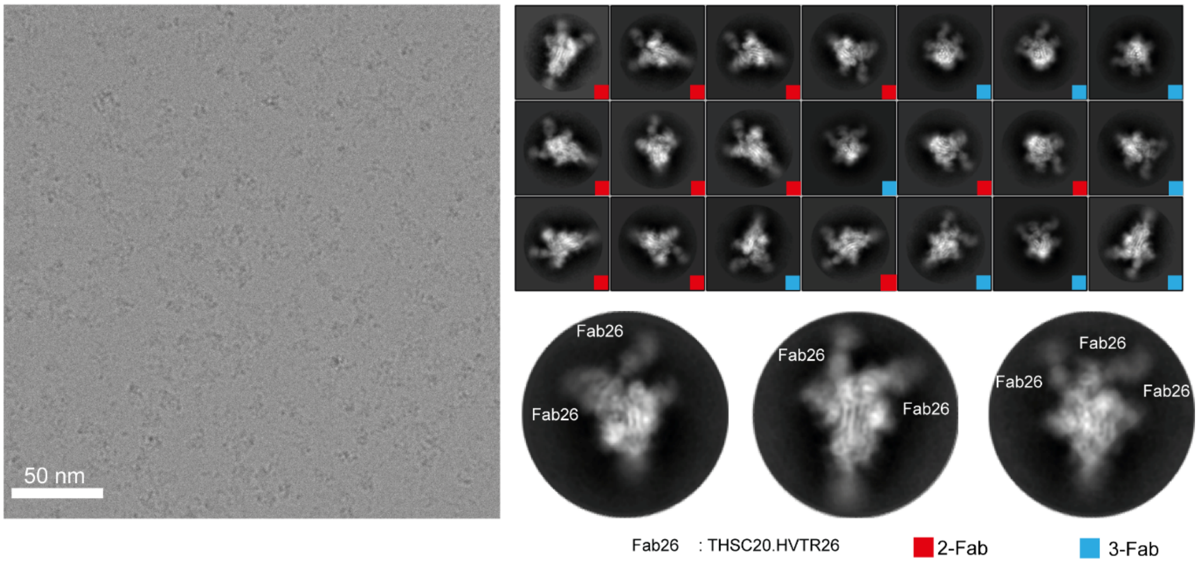


Figure S3: Cryo-electron microscopy micrograph and 2D class averages of S protein with A. Fab4 and B. Fab26 complexes

Supplementary figure 4

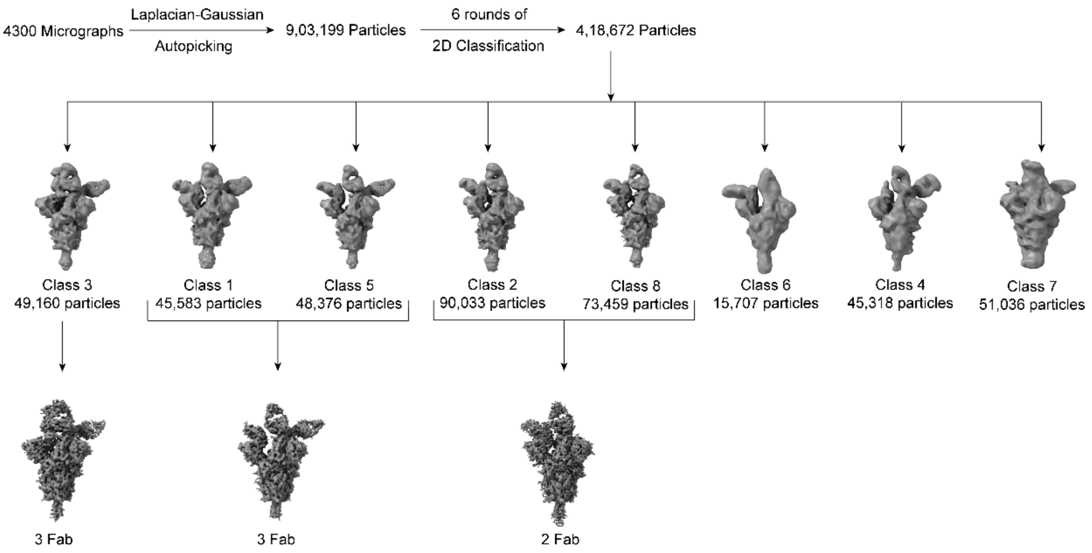


Figure S4: Workflow for the single-particle analysis (SPA) of cryo-EM for S protein: Fab4 complex: Pipeline illustrates the cryo-EM data processing, different conformations obtained and its refined structures.

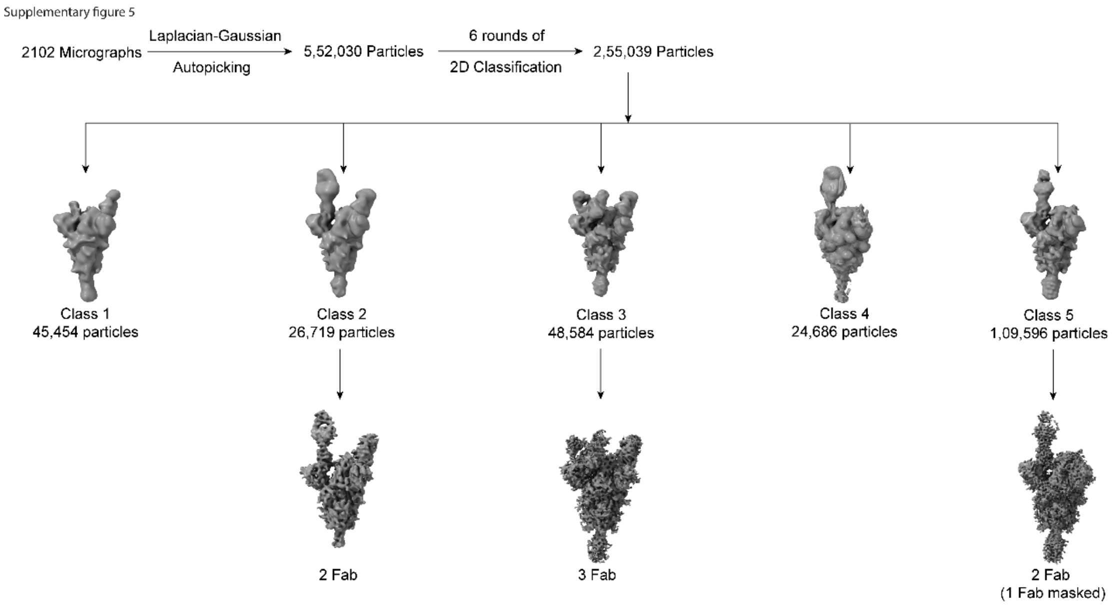


Figure S5: Workflow for the single-particle analysis (SPA) of cryo-EM for S protein: Fab26 complex: Pipeline illustrates the cryo-EM data processing, different conformations obtained and its refined structures.

Supplementary figure 6

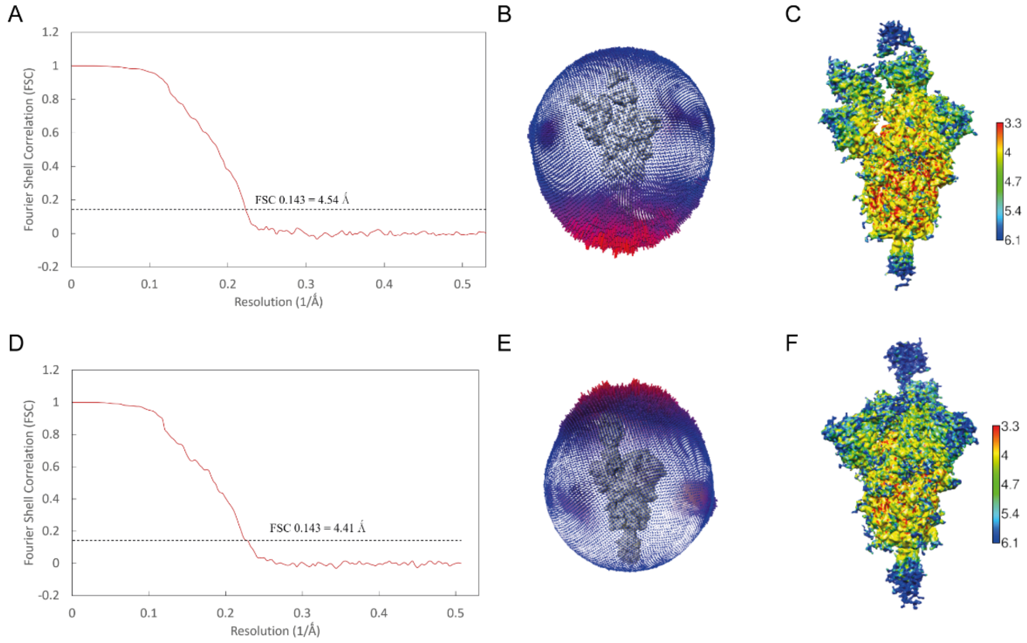


Figure S6: A, D) Gold standard Fourier shell correlation (FSC) curve calculated from the independent half maps results in 4.54 and 4.4 Å global resolution EM density map for the S protein complex with Fab4 and Fab26 respectively. B, E) Angular distribution plot for the final refined EM maps of S protein: Fab4/Fab26 complexes respectively without imposing symmetry (C1). C, F) Local resolution calculated for the high-resolution EM maps of Spike with Fab4 and Fab26 complexes respectively (C1 symmetry), coloured according to resolution calculated using ResMap.

Supplementary figure 7

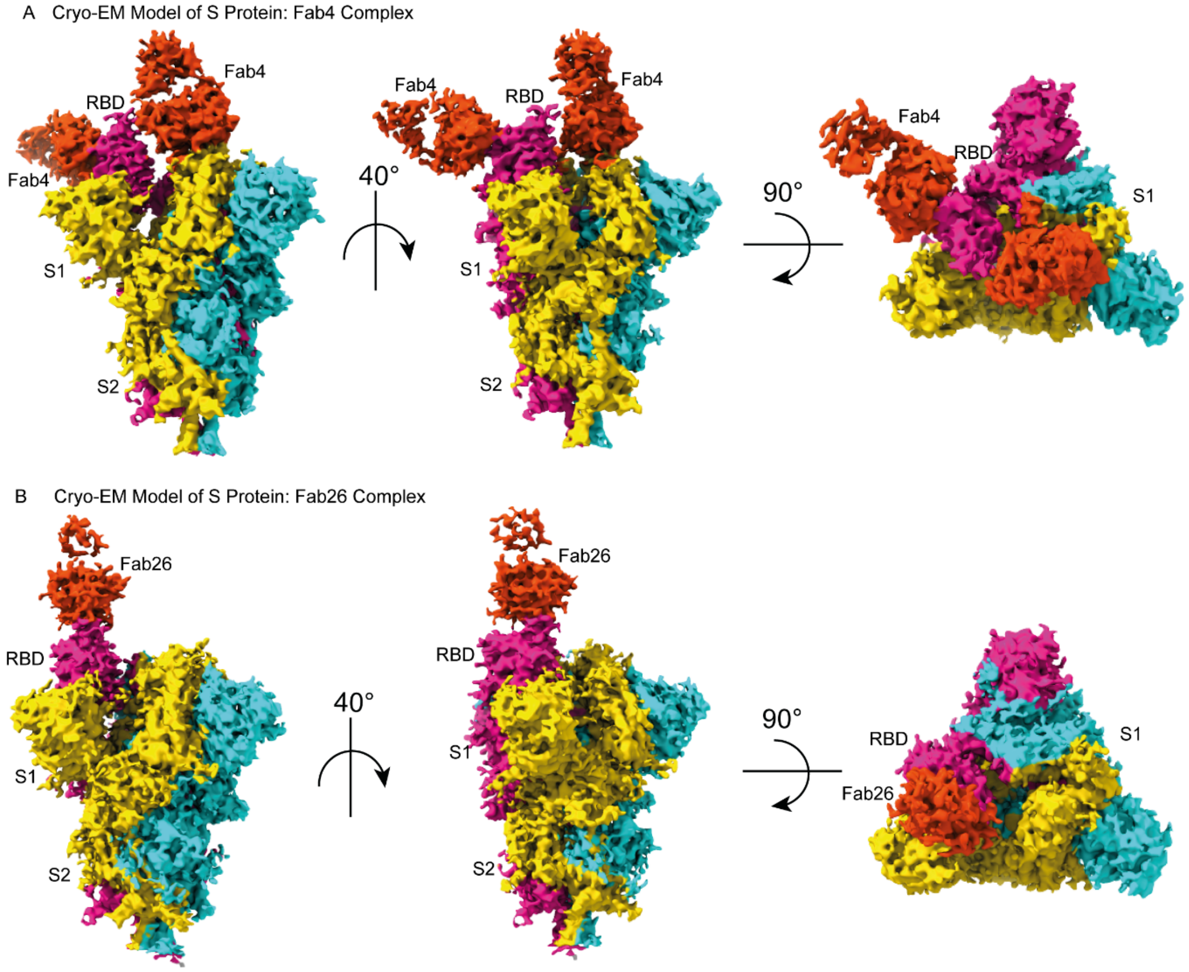


Figure S7:

Different Projections of Spike protein: Fab complex EM maps: A) High-resolution EM map showing two RBD's of S trimer bound to two Fab4's independently in upright conformation. Top and side views of the complexes were shown. B) High-resolution EM map 1-up RBD of S trimer bound to one Fab and its top and side views of the complexes were shown. Spike protomers (Pro1, 2, 3) are coloured gold, magenta, and cyan respectively, while the Fab is coloured in orange-red.
